# Neo-formation of chromosomes in bacteria

**DOI:** 10.1101/264945

**Authors:** Olivier B. Poirion, Bénédicte Lafay

## Abstract

Although the bacterial secondary chromosomes/megaplasmids/chromids, first noticed about forty years ago, are commonly held to originate from stabilized plasmids, their true nature and definition are yet to be resolved. On the premise that the integration of a replicon within the cell cycle is key to deciphering its essential nature, we show that the content in genes involved in the replication, partition and segregation of the replicons and in the cell cycle discriminates the bacterial replicons into chromosomes, plasmids, and another class of essential genomic elements that function as chromosomes. These latter do not derive directly from plasmids. Rather, they arise from the fission of a multi-replicon molecule corresponding to the co-integrated and rearranged ancestral chromosome and plasmid. All essential replicons in a distributed genome are thus neochromosomes. Having a distributed genome appears to extend and accelerate the exploration of the bacterial genome evolutionary landscape, producing complex regulation and leading to novel eco-phenotypes and species diversification.

## INTRODUCTION

Chromosomes are the only components of the genome that encode the necessary information for replication and life of the cell/organism under normal growth conditions. Their number varies across taxa, a single chromosome being the standard in bacteria (Krawiec and Riley, 1990). Evidence accumulated over the past forty years is proving otherwise: bacterial genomes can be distributed on more than one chromosome-like autonomously replicating genomic element (replicon) (Casjens, 1998; diCenzo and Finan, 2017; Mackenzie et al., 2004). The largest, primary, essential replicon (ER) in a multipartite genome corresponds to a *bona fide* chromosome and the additional, secondary, ERs (SERs) are expected to derive from accessory replicons (plasmids (Lederberg, 1998)). The most popular model of SER formation posits that a plasmid acquired by a mono-chromosome progenitor bacterium is stabilized in the genome through the transfer from the chromosome of genes essential to the cell viability (diCenzo and Finan, 2017; diCenzo et al., 2013; Slater et al., 2009). The existence in SERs of plasmid-like replication and partition systems (Dubarry et al., 2006; Egan and Waldor, 2003; Livny et al., 2007; MacLellan et al., 2004, 2006; Slater et al., 2009; Yamaichi et al., 2007) as well as experimental results (diCenzo et al., 2014) support this view. Yet, the duplication and maintenance processes of SERs contrast with the typical behaviour of plasmids for which both the timing of replication initiation and the centromere movement are random (Million-Weaver and Camps, 2014; Reyes-Lamothe et al., 2014). Indeed, the SERs share many characteristic features with chromosomes: enrichment in Dam methylation sites of the replication origin (Egan and Waldor, 2003; Gerding et al., 2015), presence of initiator titration sites (Egan and Waldor, 2003; Venkova-Canova and Chattoraj, 2011), synchronization of the replication with the cell cycle (De Nisco et al., 2014; Deghelt et al., 2014; Egan and Waldor, 2003; Egan et al., 2004; Fiebig et al., 2006; Frage et al., 2016; Kahng and Shapiro, 2003; Rasmussen et al., 2007; Srivastava et al., 2006; Stokke et al., 2011), KOPS-guided FtsK translocation (Val et al., 2008), FtsK-dependent dimer resolution system (Val et al., 2008), MatP/matS macrodomain organisation system (Demarre et al., 2014), and similar fine-scale segregation dynamics (Fiebig et al., 2006; Frage et al., 2016). Within a multipartite genome, the replication of the chromosome and that of the SER(s) are initiated at different time points (De Nisco et al., 2014; Deghelt et al., 2014; Fiebig et al., 2006; Frage et al., 2016; Rasmussen et al., 2007; Srivastava et al., 2006; Stokke et al., 2011), and use replicon-specific systems (Drevinek et al., 2008; Egan and Waldor, 2003; Galardini et al., 2013; MacLellan et al., 2004, 2006; Slater et al., 2009). Yet, they are coordinated, hence maintaining the genome stoichiometry (Deghelt et al., 2014; Egan et al., 2004; Fiebig et al., 2006; Frage et al., 2016; Stokke et al., 2011). In the few species where this was studied, the replication of the SER is initiated after that of the chromosome (De Nisco et al., 2014; Deghelt et al., 2014; Fiebig et al., 2006; Frage et al., 2016; Rasmussen et al., 2007; Srivastava, 2006; Stokke et al., 2011) under various modalities. In the Vibrionaceae, the replication of a short region of the chromosome licenses the SER duplication (Baek and Chattoraj, 2014; Kemter et al., 2018), and the advancement of the SER replication and segregation triggers the divisome assembly (Galli et al., 2016). In turn, the altering of the chromosome replication does not affect the replication initiation control of the SER in α-proteobacterium *Ensifer/Sinorhizobium meliloti* (Frage et al., 2016).

Beside the exploration of the replication/segregation mechanistic, studies of multipartite genomes, targeting a single bacterial species or genus (diCenzo et al., 2013, 2014; Dubarry et al., 2006; Mackenzie et al., 2004; Slater et al., 2009) or using a more extensive set of taxa (diCenzo and Finan, 2017; Harrison et al., 2010), relied on inadequate (replicon size, nucleotide composition, coding of core essential genes for growth and survival (diCenzo and Finan, 2017; Harrison et al., 2010; Liu et al., 2015); Figure 1) and/or oriented (presence of plasmid-type systems for genome maintenance and replication initiation (Harrison et al., 2010)) criteria to characterize the SERs.

**Figure 1.**
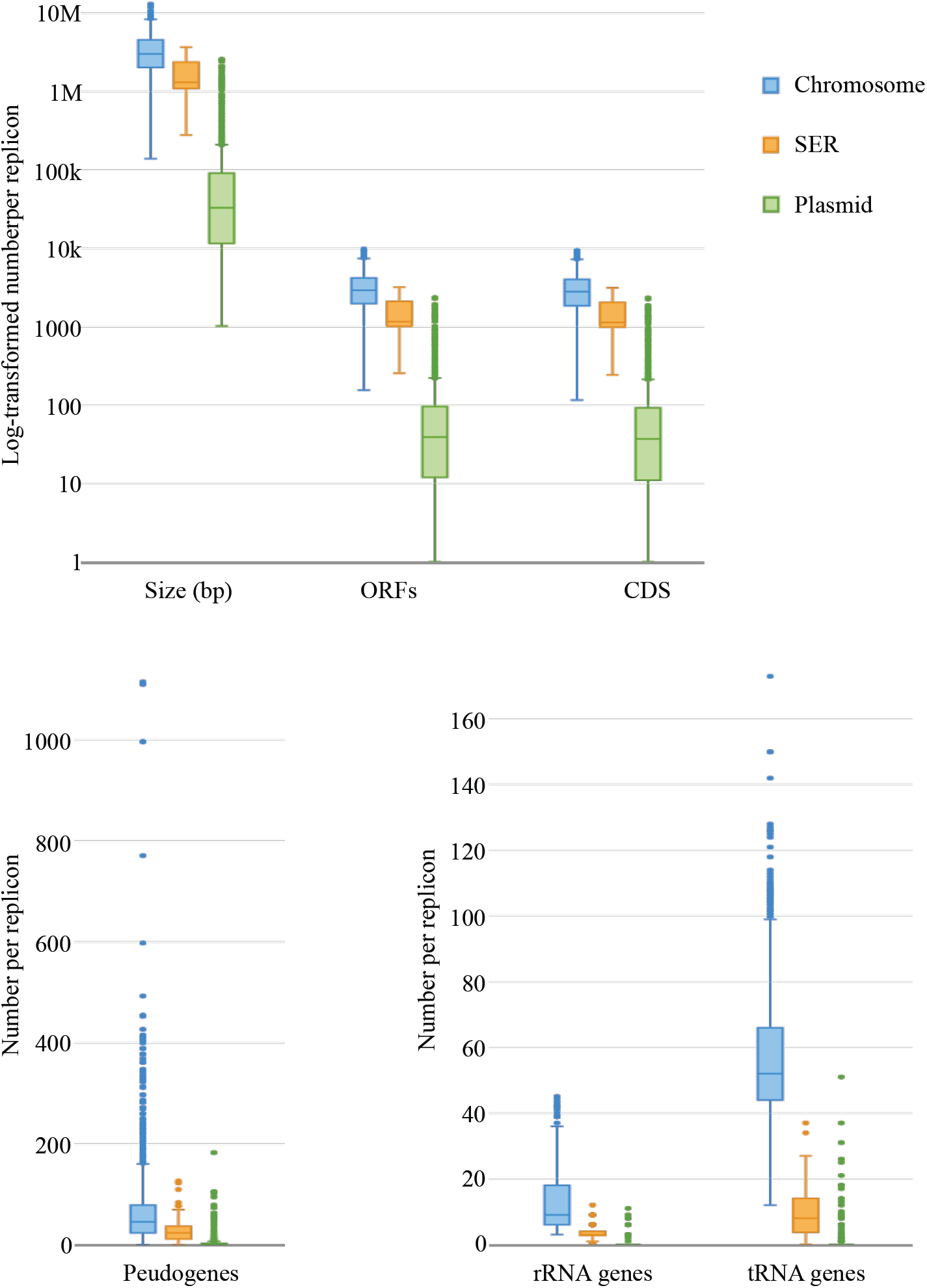
Structural features of the replicons. Boxplots of the lengths (base pairs) and numbers of genes (ORFs), protein-coding genes (CDS), pseudogenes, ribosomal RNA genes and transfer RNA genes for the 2016 chromosomes (blue), 129 SERs (orange), and 2783 plasmids (green) included in the final dataset (4928 replicons).

While clarifying the functional and evolutionary contributions of each type of replicon to a multipartite genome in given bacterial lineages (Galardini et al., 2013; Harrison et al., 2010; MacLellan et al., 2004; Slater et al., 2009), these studies produced no absolute definition of SERs (diCenzo and Finan, 2017; Harrison et al., 2010) or universal model for their emergence (diCenzo and Finan, 2017; diCenzo et al., 2013, 2014; Galardini et al., 2013; Harrison et al., 2010). We thus set out investigating the nature(s) and origin(s) of these replicons using as few assumptions as possible.

## RESULTS

### Replicon inheritance systems as diagnostic features

We did not limit our study to a particular multipartite genome or a unique gene family. Rather, we performed a global analysis encompassing all bacterial replicons whose complete sequence was available in public sequence databases (Figure 2). We reasoned that the key property discriminating the chromosomes from the plasmids is their transmission from mother to daughter cells during the bacterial cell cycle. The functions involved in the replication, partition and maintenance of a replicon, *i.e.,* its inheritance systems (ISs), thence are expected to reflect the replicon degree of integration into the host cycle.

**Figure 2.**
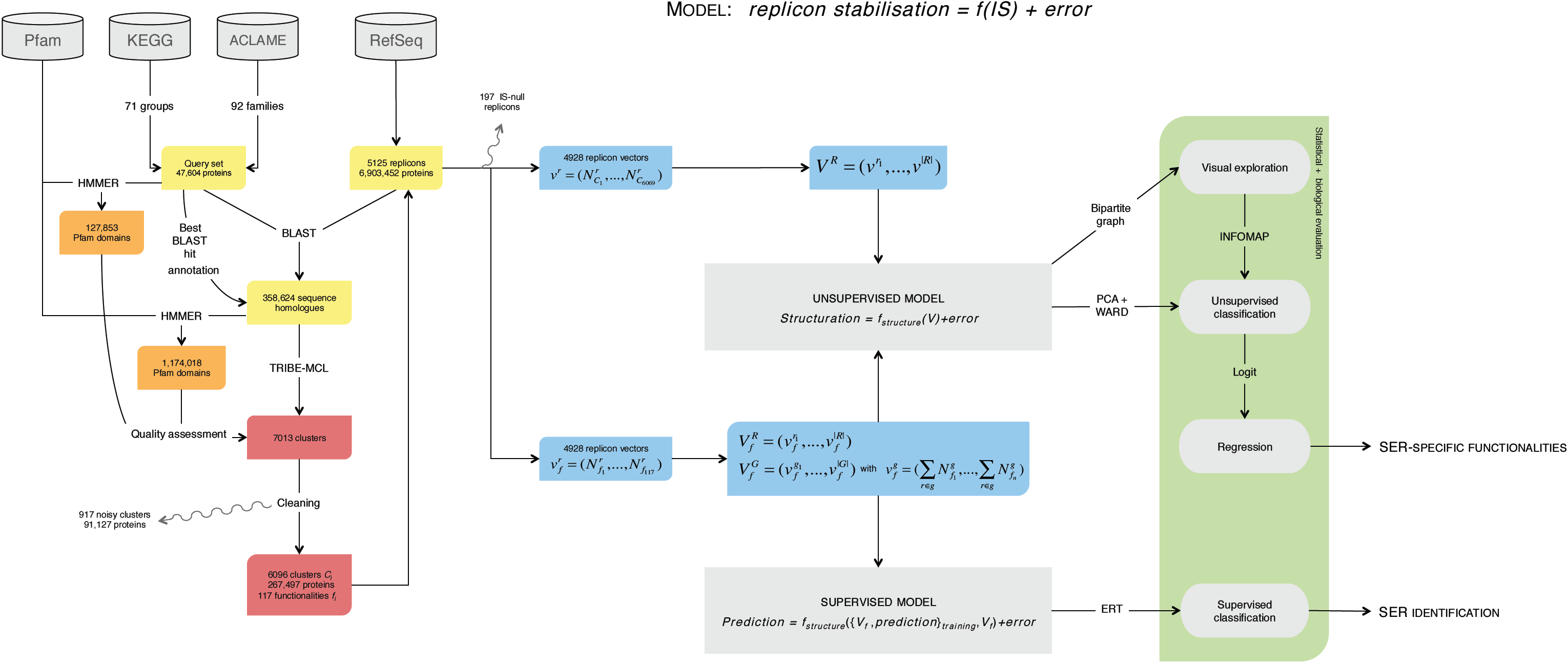
Analytical procedure.

We first faced the challenge of identifying all IS functional homologues. The inheritance of genetic information requires functionally diverse and heterogeneous actuators depending on the replicon type and the characteristics of the organism. Also, selecting sequence orthologues whilst avoiding false positives (*e.g.*, sequence paralogues) can be tricky since remote sequence homology most likely prevails among chromosome/plasmid protein-homologue pairs.

Starting from an initial dataset of 5125 replicons, we identified 358,624 putative IS functional homologues, overall corresponding to 1711 Pfam functional domains (Figure 3a), using a query set of 47,604 chromosomal and plasmidic IS-related proteins selected from the KEGG and ACLAME databases (Tables 1 and 2).

**Table 1.**
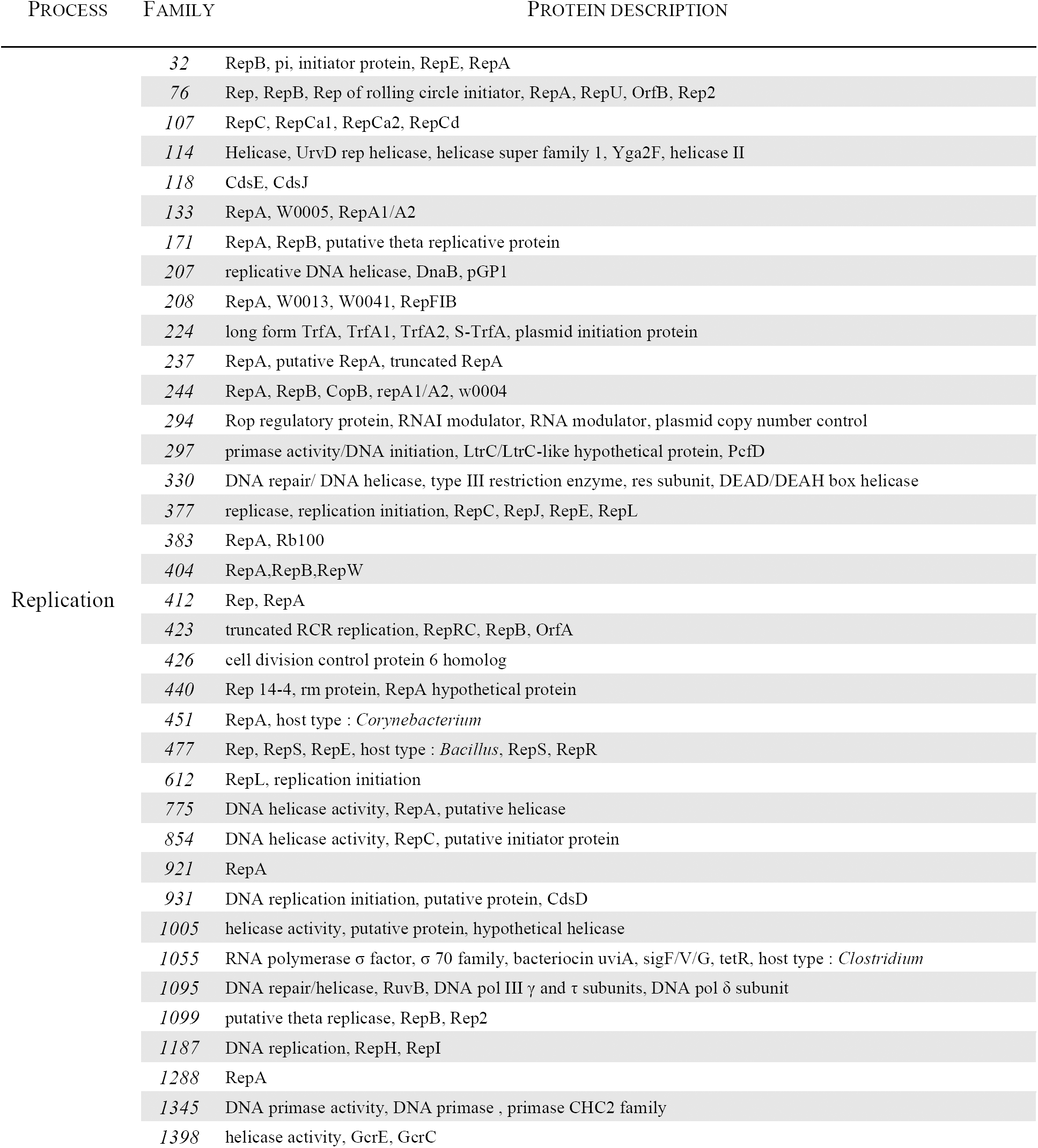
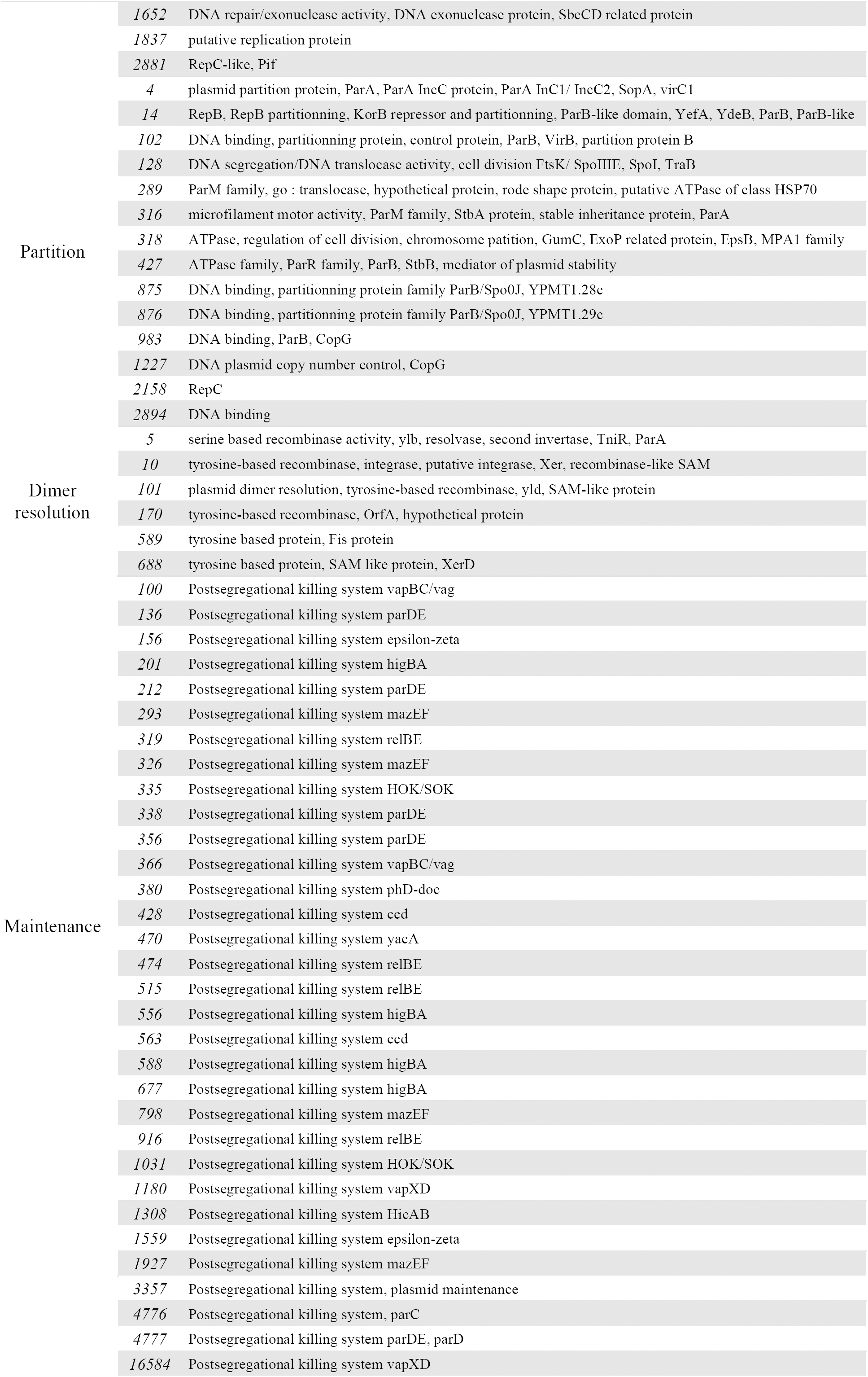
ACLAME families used in the building of the query set.

| PROCESS | FAMILY | PROTEIN DESCRIPTION |
| --- | --- | --- |
| Replication | 32 | RepB, pi, initiator protein, RepE, RepA |
|  | 76 | Rep, RepB, Rep of rolling circle initiator, RepA, RepU, OrfB, Rep2 |
|  | 107 | RepC, RepCa1, RepCa2, RepCd |
|  | 114 | Helicase, UrvD rep helicase, helicase super family 1, Yga2F, helicase II |
|  | 118 | CdsE, CdsJ |
|  | 133 | RepA, W0005, RepA1/A2 |
|  | 171 | RepA, RepB, putative theta replicative protein |
|  | 207 | replicative DNA helicase, DnaB, pGP1 |
|  | 208 | RepA, W0013, W0041, RepFIB |
|  | 224 | long form TrfA, TrfA1, TrfA2, S-TrfA, plasmid initiation protein |
|  | 237 | RepA, putative RepA, truncated RepA |
|  | 244 | RepA, RepB, CopB, repA1/A2, w0004 |
|  | 294 | Rop regulatory protein, RNAI modulator, RNA modulator, plasmid copy number control |
|  | 297 | primase activity/DNA initiation, LtrC/LtrC-like hypothetical protein, PcfD |
|  | 330 | DNA repair/ DNA helicase, type III restriction enzyme, res subunit, DEAD/DEAH box helicase |
|  | 377 | replicase, replication initiation, RepC, RepJ, RepE, RepL |
|  | 383 | RepA, Rb100 |
|  | 404 | RepA, RepB, RepW |
|  | 412 | Rep, RepA |
|  | 423 | truncated RCR replication, RepRC, RepB, OrfA |
|  | 426 | cell division control protein 6 homolog |
|  | 440 | Rep 14-4, rm protein, RepA hypothetical protein |
|  | 451 | RepA, host type : *Corynebacterium* |
|  | 477 | Rep, RepS, RepE, host type : *Bacillus*, RepS, RepR |
|  | 612 | RepL, replication initiation |
|  | 775 | DNA helicase activity, RepA, putative helicase |
|  | 854 | DNA helicase activity, RepC, putative initiator protein |
|  | 921 | RepA |
|  | 931 | DNA replication initiation, putative protein, CdsD |
|  | 1005 | helicase activity, putative protein, hypothetical helicase |
| | 1055 | RNA polymerase $\sigma$ factor, $\sigma$ 70 family, bacteriocin uviA, sigF/V/G, tetR, host type : *Clostridium* |
| | 1095 | DNA repair/helicase, RuvB, DNA pol III $\gamma$ and $\tau$ subunits, DNA pol $\delta$ subunit |
|  | 1099 | putative theta replicase, RepB, Rep2 |
|  | 1187 | DNA replication, RepH, RepI |
|  | 1288 | RepA |
|  | 1345 | DNA primase activity, DNA primase , primase CHC2 family |
|  | 1398 | helicase activity, GcrE, GcrC |
|  | 1652 | DNA repair/exonuclease activity, DNA exonuclease protein, SbcCD related protein |
|  | 1837 | putative replication protein |
|  | 2881 | RepC-like, Pif |
| Partition | 4 | plasmid partition protein, ParA, ParA IncC protein, ParA Inc1/ IncC2, SopA, virC1 |
|  | 14 | RepB, RepB partitioning, KorB repressor and partitioning, ParB-like domain, YefA, YdeB, ParB, ParB-like |
|  | 102 | DNA binding, partitioning protein, control protein, ParB, VirB, partition protein B |
|  | 128 | DNA segregation/DNA translocase activity, cell division FtsK/ SpoIIIE, SpoI, TraB |
|  | 289 | ParM family, go : translocase, hypothetical protein, rode shape protein, putative ATPase of class HSP70 |
|  | 316 | microfilament motor activity, ParM family, StbA protein, stable inheritance protein, ParA |
|  | 318 | ATPase, regulation of cell division, chromosome partition, GumC, ExoP related protein, EpsB, MPA1 family |
|  | 427 | ATPase family, ParR family, ParB, StbB, mediator of plasmid stability |
|  | 875 | DNA binding, partitioning protein family ParB/Spo0J, YPMT1.28c |
|  | 876 | DNA binding, partitioning protein family ParB/Spo0J, YPMT1.29c |
|  | 983 | DNA binding, ParB, CopG |
|  | 1227 | DNA plasmid copy number control, CopG |
|  | 2158 | RepC |
|  | 2894 | DNA binding |
| Dimer resolution | 5 | serine based recombinase activity, ylb, resolvase, second invertase, TniR, ParA |
|  | 10 | tyrosine-based recombinase, integrase, putative integrase, Xer, recombinase-like SAM |
|  | 101 | plasmid dimer resolution, tyrosine-based recombinase, yld, SAM-like protein |
|  | 170 | tyrosine-based recombinase, OrfA, hypothetical protein |
|  | 589 | tyrosine based protein, Fis protein |
|  | 688 | tyrosine based protein, SAM like protein, XerD |
| Maintenance | 100 | Postsegregational killing system vapBC/vag |
|  | 136 | Postsegregational killing system parDE |
|  | 156 | Postsegregational killing system epsilon-zeta |
|  | 201 | Postsegregational killing system higBA |
|  | 212 | Postsegregational killing system parDE |
|  | 293 | Postsegregational killing system mazEF |
|  | 319 | Postsegregational killing system relBE |
|  | 326 | Postsegregational killing system mazEF |
|  | 335 | Postsegregational killing system HOK/SOK |
|  | 338 | Postsegregational killing system parDE |
|  | 356 | Postsegregational killing system parDE |
|  | 366 | Postsegregational killing system vapBC/vag |
|  | 380 | Postsegregational killing system phD-doc |
|  | 428 | Postsegregational killing system ccd |
|  | 470 | Postsegregational killing system yacA |
|  | 474 | Postsegregational killing system relBE |
|  | 515 | Postsegregational killing system relBE |
|  | 556 | Postsegregational killing system higBA |
|  | 563 | Postsegregational killing system ccd |
|  | 588 | Postsegregational killing system higBA |
|  | 677 | Postsegregational killing system higBA |
|  | 798 | Postsegregational killing system mazEF |
|  | 916 | Postsegregational killing system relBE |
|  | 1031 | Postsegregational killing system HOK/SOK |
|  | 1180 | Postsegregational killing system vapXD |
|  | 1308 | Postsegregational killing system HicAB |
|  | 1559 | Postsegregational killing system epsilon-zeta |
|  | 1927 | Postsegregational killing system mazEF |

|  |  |
| --- | --- |
| 3357 | Postsegregational killing system, plasmid maintenance |
| 4776 | Postsegregational killing system, parC |
| 4777 | Postsegregational killing system parDE, parD |
| 16584 | Postsegregational killing system vapXD |

**Table 2.**
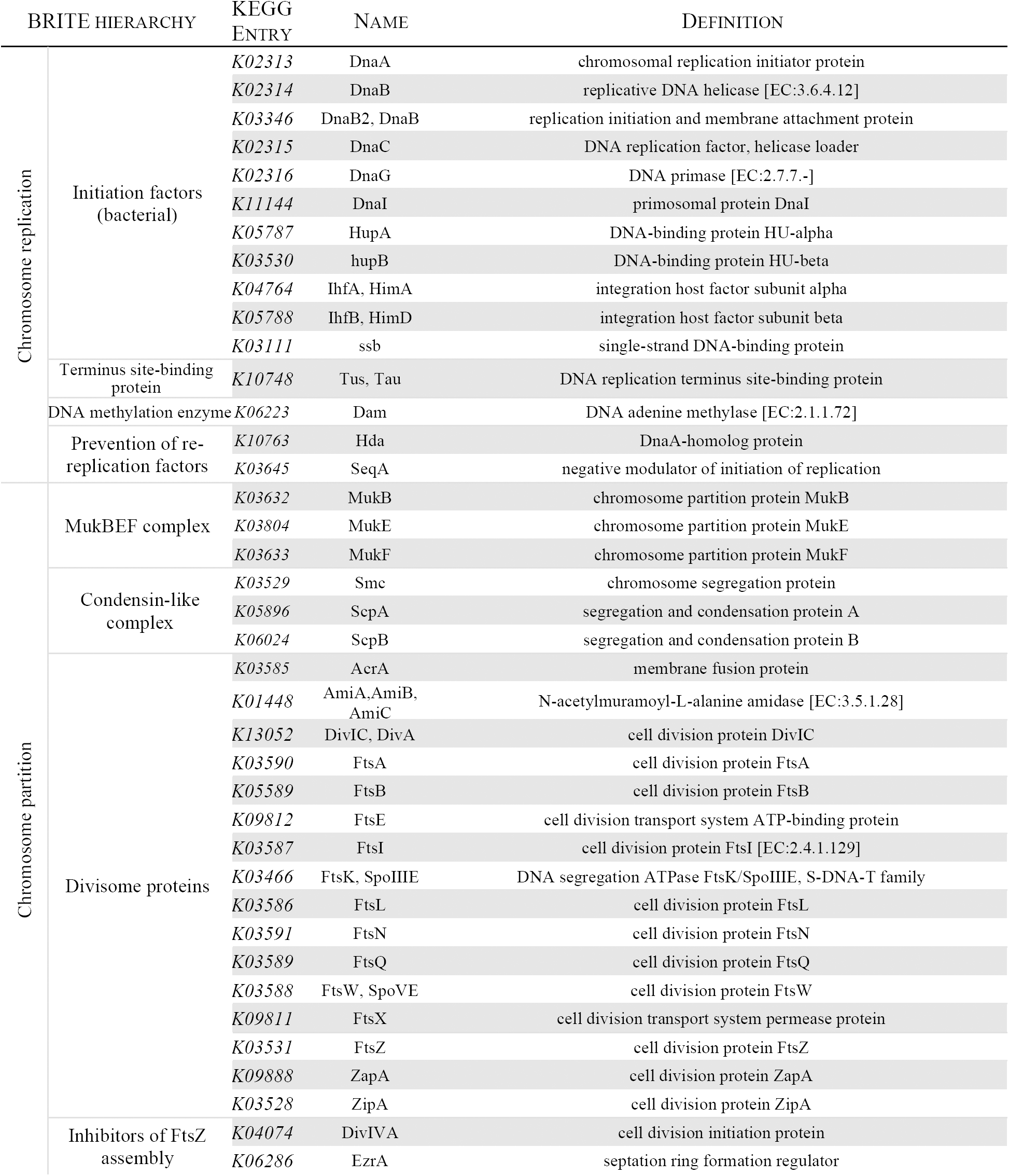
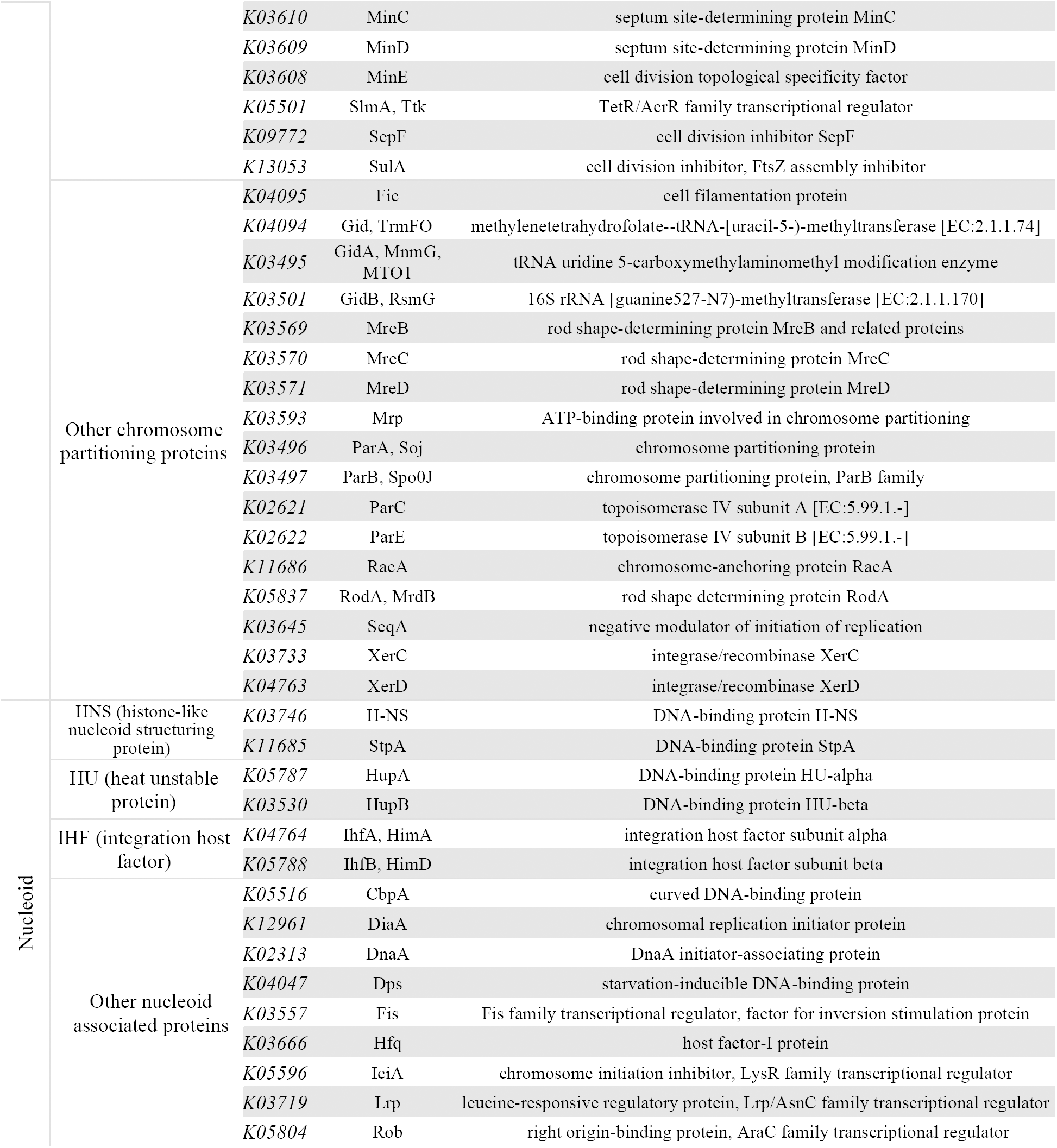
KEGG “Prokaryotic-type chromosome” orthology groups used in the building of the query set.

**Figure 3.**
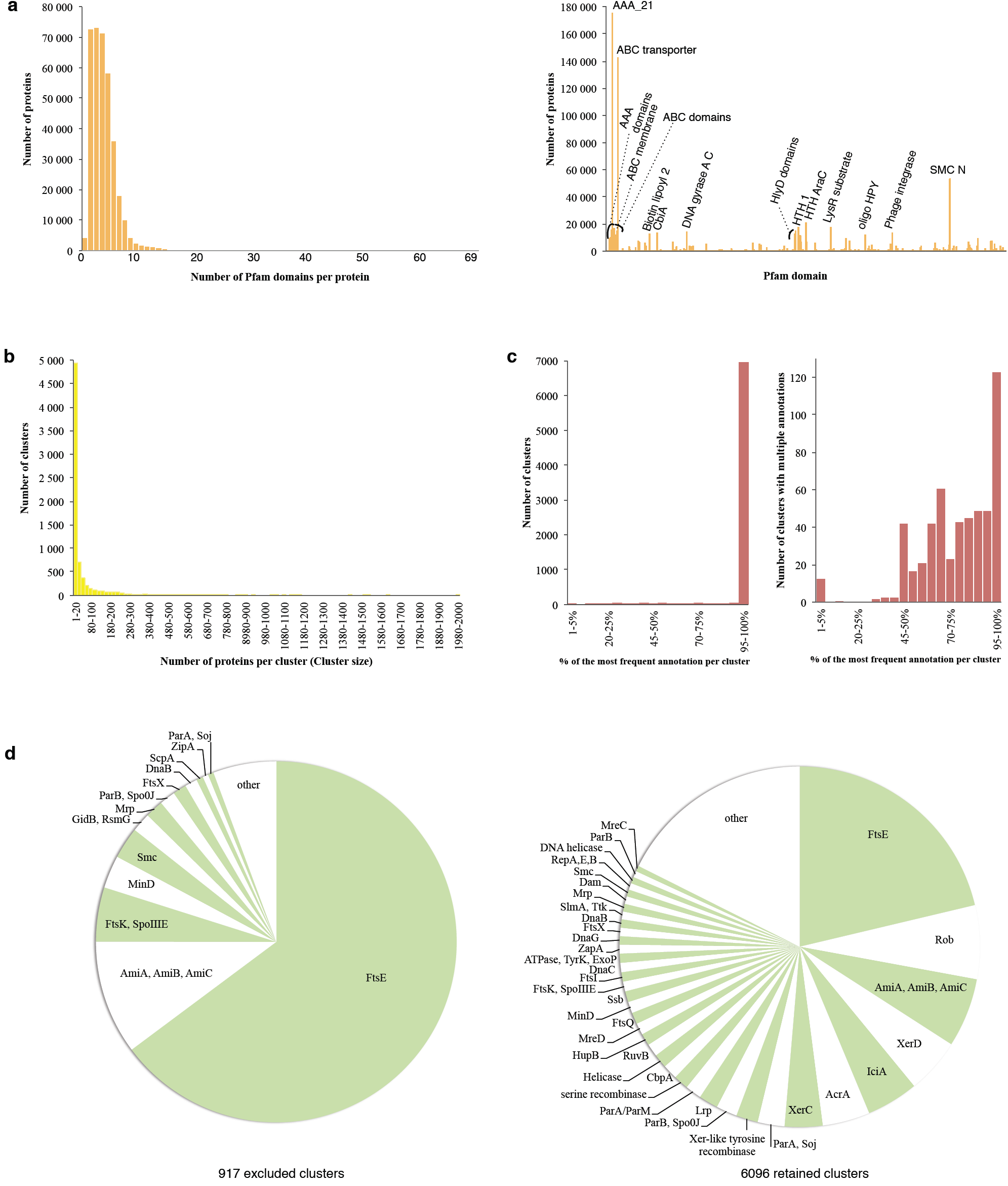
Properties of the IS clustering. (a) Frequency distribution of the 358,624 putative IS protein homologues according to their number of functional domains (0 to 39) per protein (left), and occurrences of the 1711 functional Pfam domains (right). The 20 top most frequently encountered functional domains are indicated, (b) Size distribution of the 7013 clusters, each comprising from a single to 1990 proteins, (c) Percentage distribution of the most frequent annotation per cluster among all clusters (left) and among clusters with multiple innotations (right), (d) Distribution of the most frequent annotation per cluster among the 917 excluded clusters (left) and the 3096 clusters retained for the analysis (right).

We then inferred 7013 homology groups using a clustering procedure and named the clusters after the most frequent annotation found among their proteins (Figure 3b,c). Most clusters were characterized by a single annotation whilst the remaining few (4.7%) each harbored from 2 to 710 annotations, the most frequent annotation in a cluster generally representing more than half of all annotations (Figure 3c). The removal of false positives left 267,497 IS protein homologues distributed in 6096 clusters (Figure 3d) and coded by 4928 replicons out of the initial replicon dataset. Following the Genbank/RefSeq annotations, our final dataset comprised 2016 complete genome sets corresponding to 3592 replicons (2016 chromosomes, 129 SERs, and 1447 plasmids) and 1336 plasmid genomes (Supplementary table 1), irregularly distributed across the bacterial phylogeny (Figure 4a). Multi-ER genomes are observed in 5.0% of all represented bacterial genera and constitute 5.7% of the complete genomes (averaged over genera) available at the time of study (Figure 4b). They are merely incidental (0.2% in Firmicutes) or reach up to almost one third of the genomes (30.1% in β-Proteobacteria) depending on the lineage, and are yet to be observed in most bacterial phyla, possibly because of the poor representation of some lineages. Although found in ten phyla, they occur more than once per genus in only three of them: Bacteroidetes, Proteobacteria and Spirochaetae.

**Figure 4.**
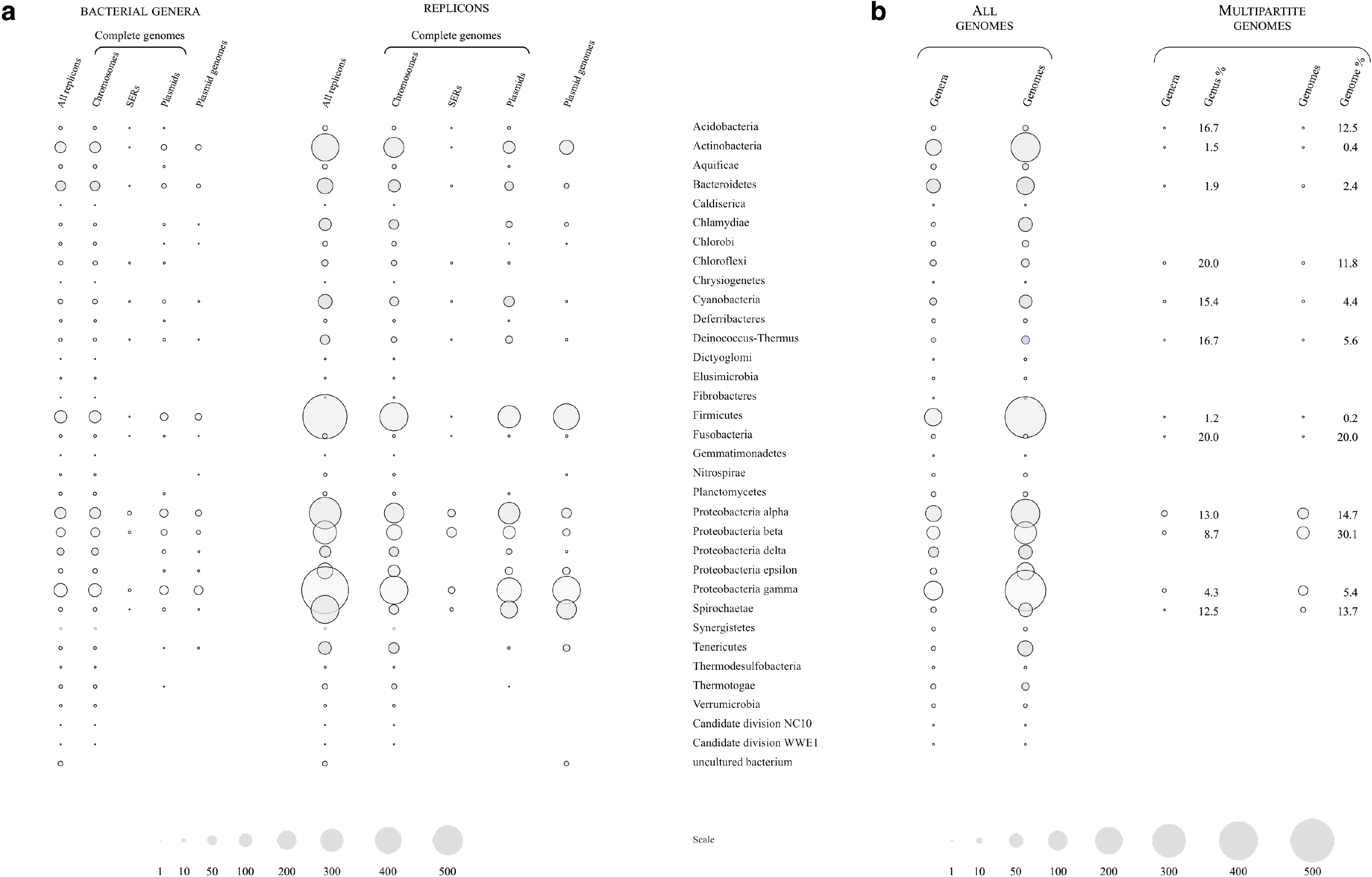
Taxonomic structure of the replicon dataset. Numbers of replicons (a) and complete genomes (b), and represented bacterial genera are shown according to datasets and host taxonomy. Surfaces represent the numbers of bacterial genera, replicons or genomes within each category. Percentages of multipartite genomes and corresponding bacterial genera are calculated for each host phylum or class (Proteobacteria).

### Exploration of the replicon diversity

We explored the differences and similarities of the bacterial replicons with regard to their IS usage using a data mining and machine learning approach (Methods). The 6096 retained IS clusters were used as distinct variables to ascribe each of the 4928 replicons with a vector according to its IS usage profile. We transformed these data into bipartite graphs depending on the number of proteins from the IS clusters coded by each replicon. Bipartite graphs display both the vectors (replicons) and the variables (protein clusters) together with their respective connections, and allow the interactive exploration of the data. The majority of the replicons are interconnected (Figure 5) as testimony of the shared evolutionary history of their IS sequences. Chromosomes and plasmids form overall distinct groups and communities with varying degree of connectivity depending on their functional specificities (Figure 5a) as well as on the bacterial taxonomy of their hosts (Figure 5c). They nonetheless share many ISs, bearing witness to the continuity of the genomic material and the extensive exchange of genetic material within bacterial genomes. The occurrence of poorly IS cluster-connected plasmids within a group of chromosomes did not consistently reflect a true relationship and rather resulted from shared connections to a very small number (as low as one) of common ISs. While being interconnected to both chromosomes and plasmids *via* numerous IS clusters, the SERs generally stand apart from either these types of replicons and gather at the chromosome-plasmid interface (Figure 5a,b). Their IS usage is neither chromosome-like nor plasmidlike, suggesting that they may constitute a separate category of replicons. This is most tangible in the case of the proteobacterial lineages where SERs occur most frequently (top of Figure 5b).

**Figure 5.**
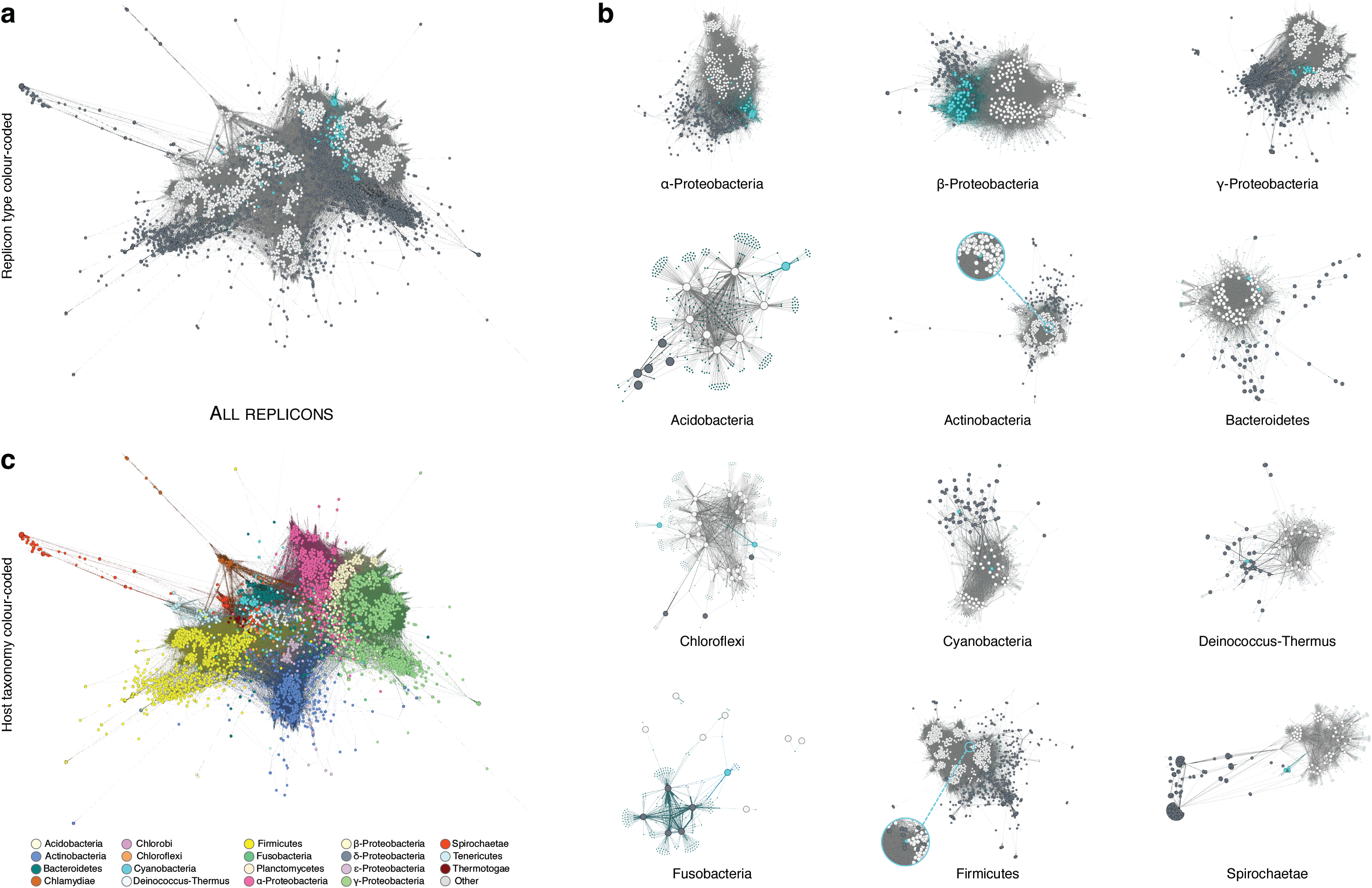
Visualisation of the replicon IS-based relationshipss. Gephi-generated bipartite-graphs for the whole dataset (a and b) or groups of replicons following the host taxonomy (c). Nodes correspond to the replicons (large dots) or the clusters of IS proteins (small dots). Edges linking replicons and protein clusters reflect the presence on a replicon of at least one protein of a protein cluster. Colouring according to replicon type (a and c): chromosomes (white), plasmids (grey), and SERs (blue), or according to host taxonomy (b).

All SERs in the β- and γ-Proteobacteria, and most in the α-Proteobacteria are linked to remarkable chromosome-type IS clusters, such as AcrA, IciA, FtsE, HN-S and Lrp, as well as to plasmid-like ParA/ParB, Rep and PSK IS clusters. A similar pattern is observed for the SERs in actinobacterium *Nocardiopsis dassonvillei,* firmicute *Butyrivibrio proteoclasticus,* and chloroflexi *Sphaerobacter thermophilus* and *Thermobaculum terrenum* (Figure 5b). Interestingly, DNA primase DnaG-annotated clusters connect the SERs present in all but one *Burkholderia* species (β-Proteobacteria) as well as the chromosomes of all other bacteria. Since the sole exception, *B. rhizoxinica,* possesses a SER-less reduced genome as an adaption to intracellular life, the *Burkholderia* SERs likely originated from a single event prior to the diversification of the genus, possibly in relation to the speciation event that gave rise to this lineage. The second SERs harbored by only some *Burkholderia* species exhibit a higher level of interconnection to plasmids, as do the SERs in α-proteobacterium *Sphingobium,* cyanobacterium *Cyanothece* sp. ATCC 51142, *Deinococcus radiodurans* (Deinococcus-Thermus) and fusobacterium *Ilyobacter polytropus.* This points to an incomplete stabilization of the SERs into the genome that may reflect a recent, ongoing, event of integration and/or differing selective pressures at play depending on the bacterial lineages. At odds with these observations, some SERs group unambiguously with chromosomes. The SERs in α-Proteobacteria *Asticcacaulis excentricus* and *Paracoccus denitrificans,* Bacteroidetes *Prevotella intermedia* and *P. melaninogenica,* acidobacterium *Chloracidobacterium thermophilum*, and cyanobacterium *Anabaena* sp. 90 bear higher levels of interconnection to chromosomes than to plasmids or other SERs. Indeed, the SERs in *Prevotella* spp. are hardly linked to plasmids, and the few plasmid-like IS proteins that *C. thermophilum* SER codes (mostly Rep, Helicase and PSK), albeit found in plasmids occurring in other phyla, are observed in none of the Acidobacteria plasmids. An extreme situation is met in *Leptospira* spp. (Spirochaetae) whose SERs are each linked to only three or four (out of a total of six) chromosome-like IS clusters, always including ParA and ParB. Interestingly, the ParA cluster appears to be specific to Spirochaetae chromosomes with the notable exception of one plasmid found in Leptospiraceae *Turneriella parva.*

### IS-based relationships of the replicons

We submitted the bipartite graph of the whole dataset to a community structure detection algorithm (INFOMAP) that performs a random walk along the edges connecting the graph vertices. We expected the replicon communities to be trapped in high-density regions of the graph. We also performed a dimension reduction by Principal Component Analysis followed by a hierarchical clustering procedure (WARD). The clustering solutions (Supplementary tables 2 and 3) were meaningful (high values reached by the stability criterion scores), and biologically relevant (efficient separation of the chromosomes from the plasmids; high *homogeneity* values) using either method (Table 3). In another experiment, we considered each genus as a unique sample and averaged the variables over the replicons of the different species for each replicon type. The aim was to control for the disparity in taxon representation of the replicons. This dataset produced overall similar albeit slightly less stable clusters (lower *homogeneity* values). Taxonomically homogeneous clusters of chromosomes were best retrieved using the coupling of dimension reduction and hierarchical clustering with a large enough number of clusters (*homogeneity* scores up to 0.93). In turn, the community detection algorithm was more efficient in recovering the underlying taxonomy of replicons (higher value of *completeness*), and was sole able to identify small and scattered plasmid clusters (Supplementary tables 2 and 3).

**Table 3.**
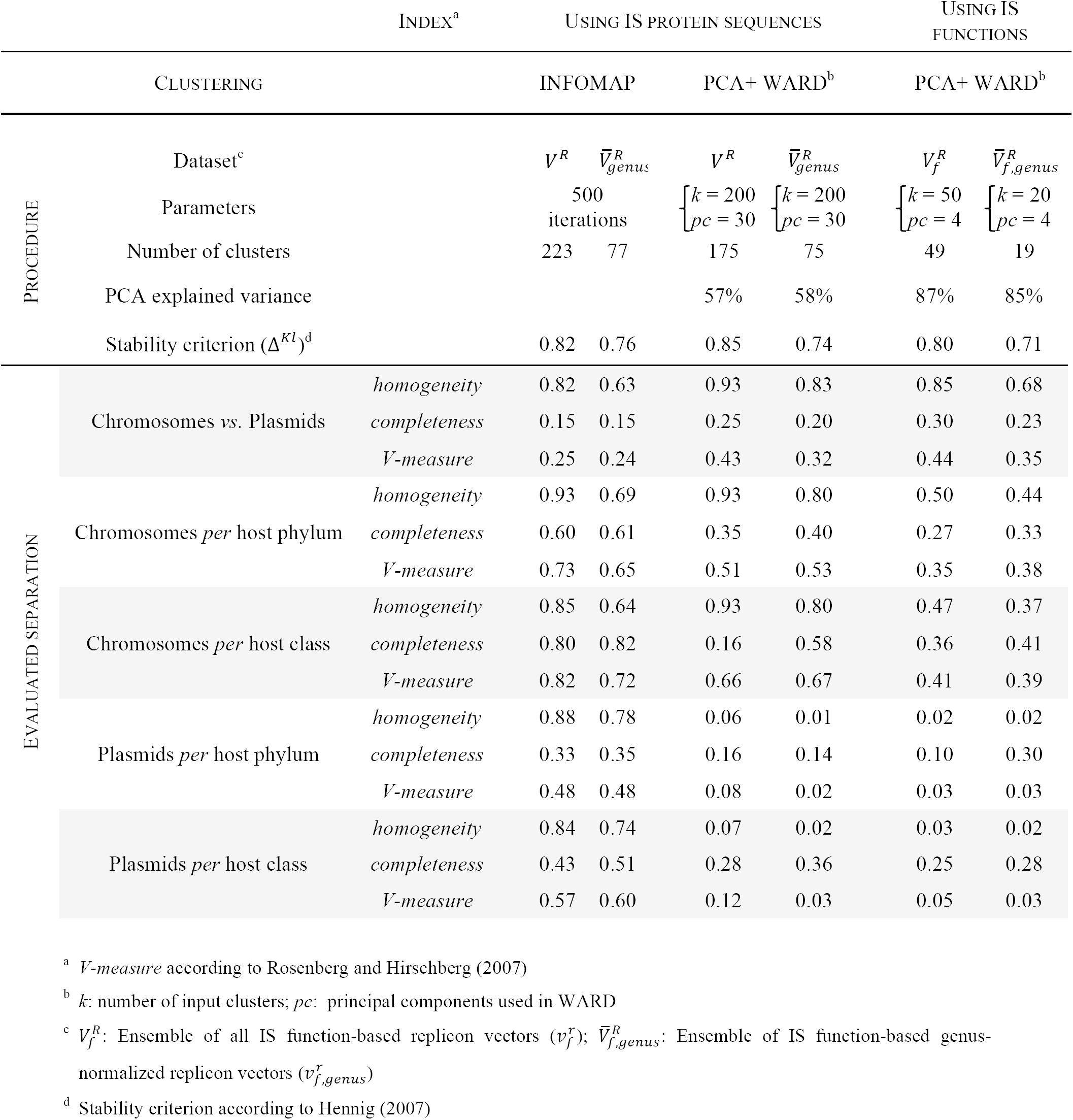
Evaluation of the replicon IS-based clusterings.

The plasmid clusters obtained using PCA+WARD lacked taxonomical patterning and, although highly stable, only reflected the small Euclidian distances existing among the plasmid replicons (*e.g*., one cluster of 2656 plasmids had a stability score of 0.975). The clusters obtained with INFOMAP mirrored the taxonomical structure of the data, suggesting that the taxonomic signal, expected to be associated to the chromosomes, is preserved among the IS protein families functionally specifying the plasmids. The presence of a majority of the SERs amongst the chromosome clusters generated by INFOMAP confirmed the affinities between these two genomic elements and the clear individuation of the SERs from the plasmids. However, the larger number of chromosomal ISs often caused the PCA+WARD approach to place SERs into plasmid clusters. The SERs in *Butyrivibrio, Deinococcus, Leptospira* and *Rhodobacter* spp. grouped consistently with plasmids while the SERs in *Vibrionaceae* and *Brucellaceae* formed specific clusters (Table 4). Burkholderiales and *Agrobacterium* SERs, whose homogenous clusters tended to be unstable, exhibited a higher affinity to plasmids overall. The SERs of *Asticaccaulis, Paracoccus* and *Prevotella* spp. associated stably with chromosomes using the two clustering methods (Table 4a,b) and possess IS profiles that set them apart from both the plasmids and the other SERs.

**Table 4.**
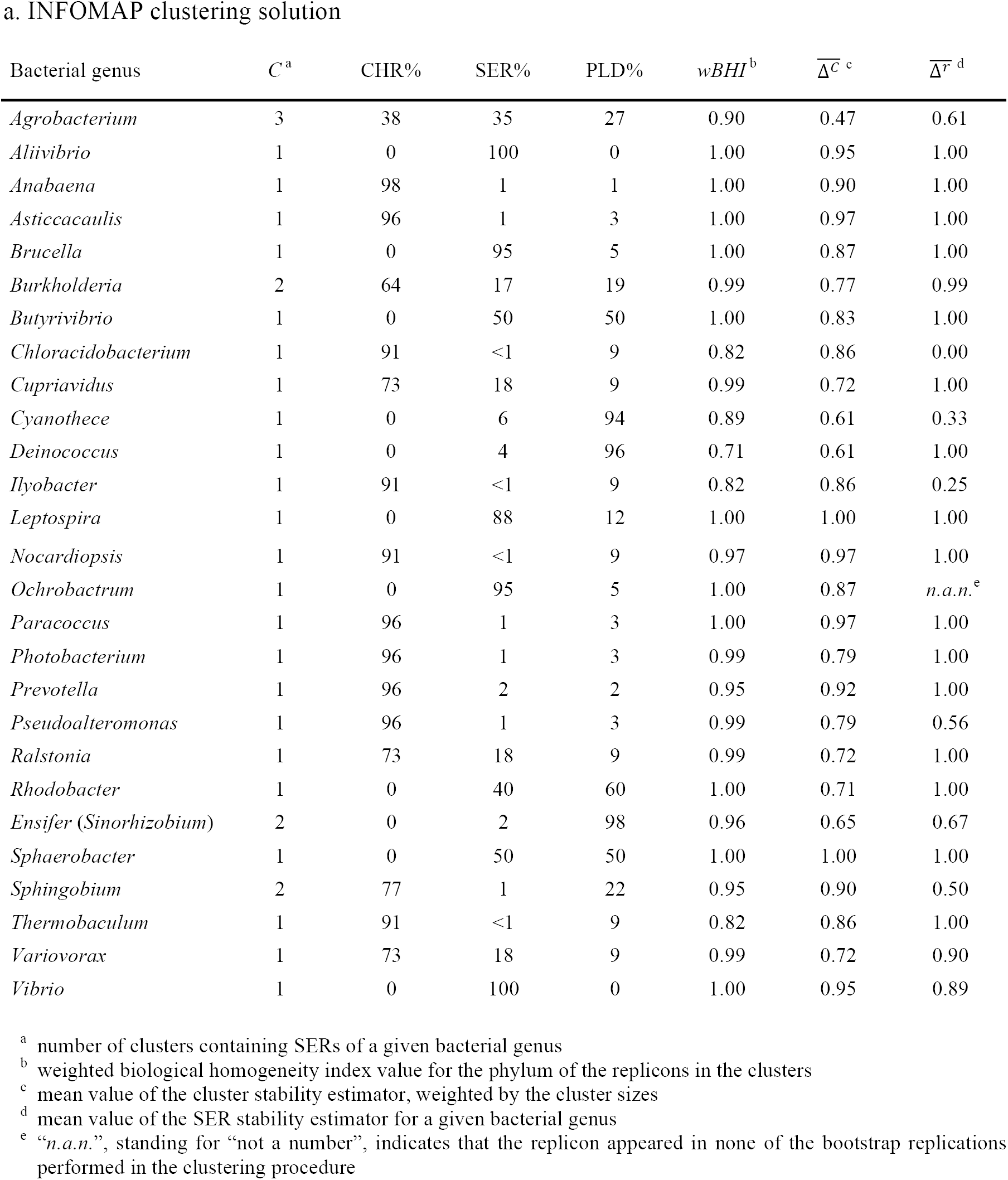
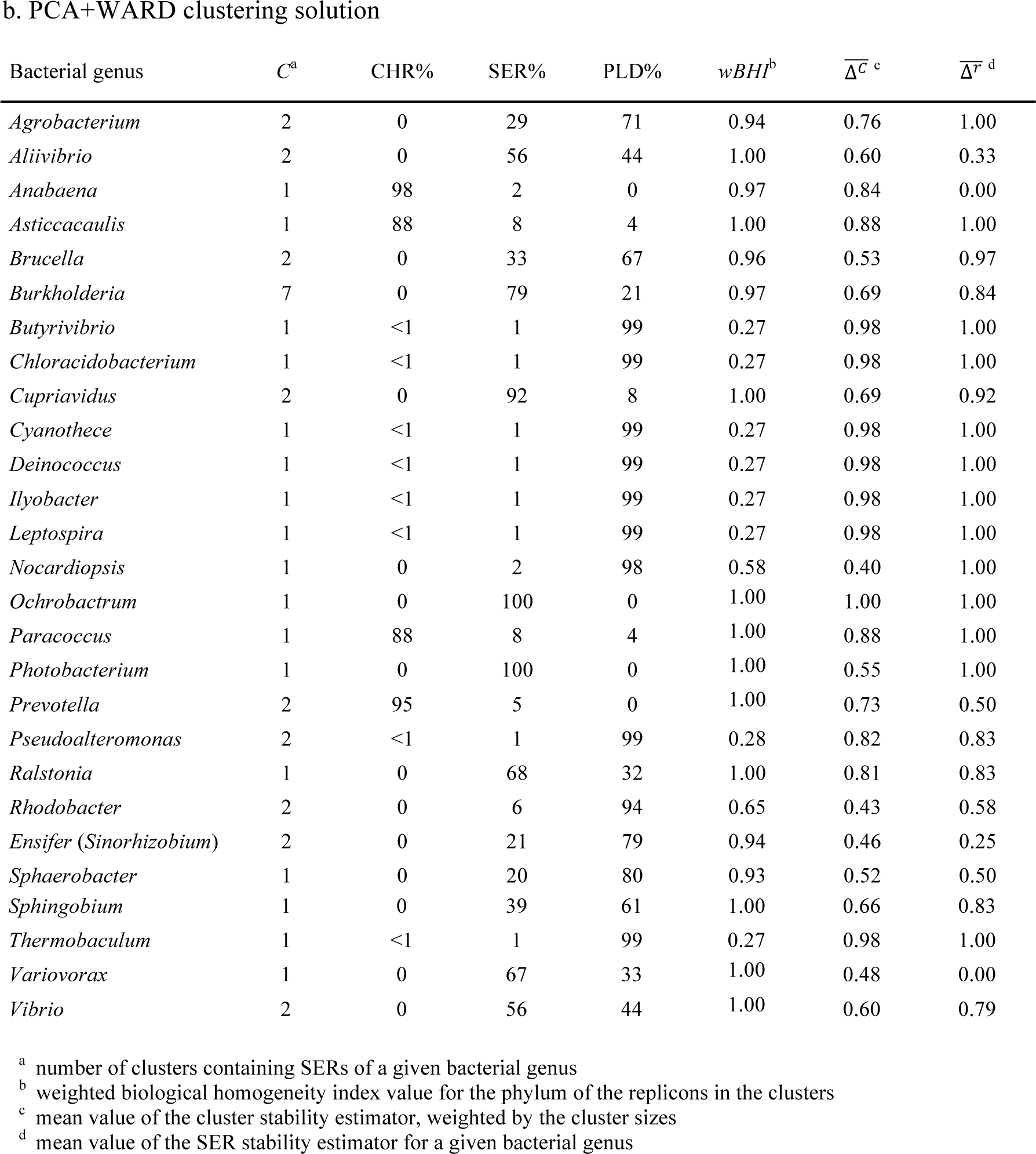
IS protein cluster-based unsupervised classification of SERs.

We reached similar conclusions when performing a PCA+WARD clustering using the 117 functional annotations of the IS protein clusters rather than the IS clusters themselves (Tables 3 and 5; Supplementary table 4).

**Table 5.**
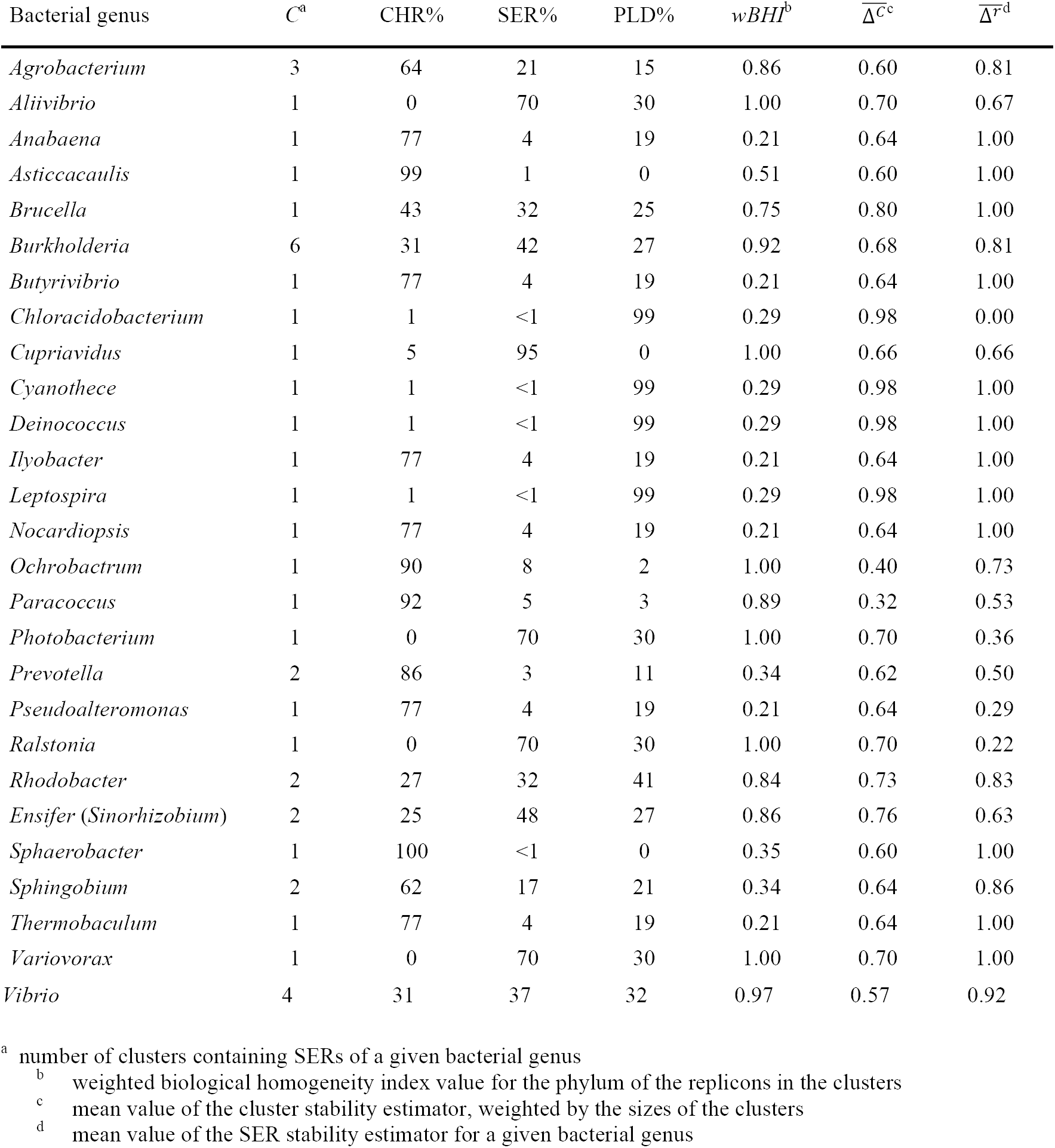
Function-based unsupervised classification of SERs using PCA+WARD.

Remarkably, in this latter analysis, the chromosomes in the multipartite genomes of *Prevotella intermedia* and *P. melaninogenica* were more similar to plasmids than to other groups of chromosomes and to single chromosomes in other *Prevotella* species (*P. denticola* and *P. ruminicola*).

### SER-specifying IS functions

Next, we searched which of the IS functions are specific to the SERs. We performed several logistic regression analyses to identify over- or under-represented ISs and to assess their respective relevance to each class of replicons. Because of their comparatively small number, all SERs were assembled into a single group despite their disparity. A hundred and one IS functionalities (96% of KEGG-annotated chromosome-like functions and 72% of ACLAME-annotated plasmid-like functions) were significantly enriched in one replicon category over the other (Table 6). The large majority of the IS functions differentiates the chromosomes from the plasmids. The latter are only determined by ISs corresponding to ACLAME annotations Rep, Rop and TrfA, involved in initiation of plasmid replication, and ParA and ParB, dedicated to plasmid partition. Some KEGG-annotated functions, *e.g.,* DnaA, DnaB or FtsZ, appear to be more highly specific to chromosomes (higher *OR* values) than others such as DnaC, FtsE or H-NS (lower *OR* values). Strikingly, very few functions distinguish significantly the chromosomes from the SERs, by contrast with plasmids.

**Table 6.**
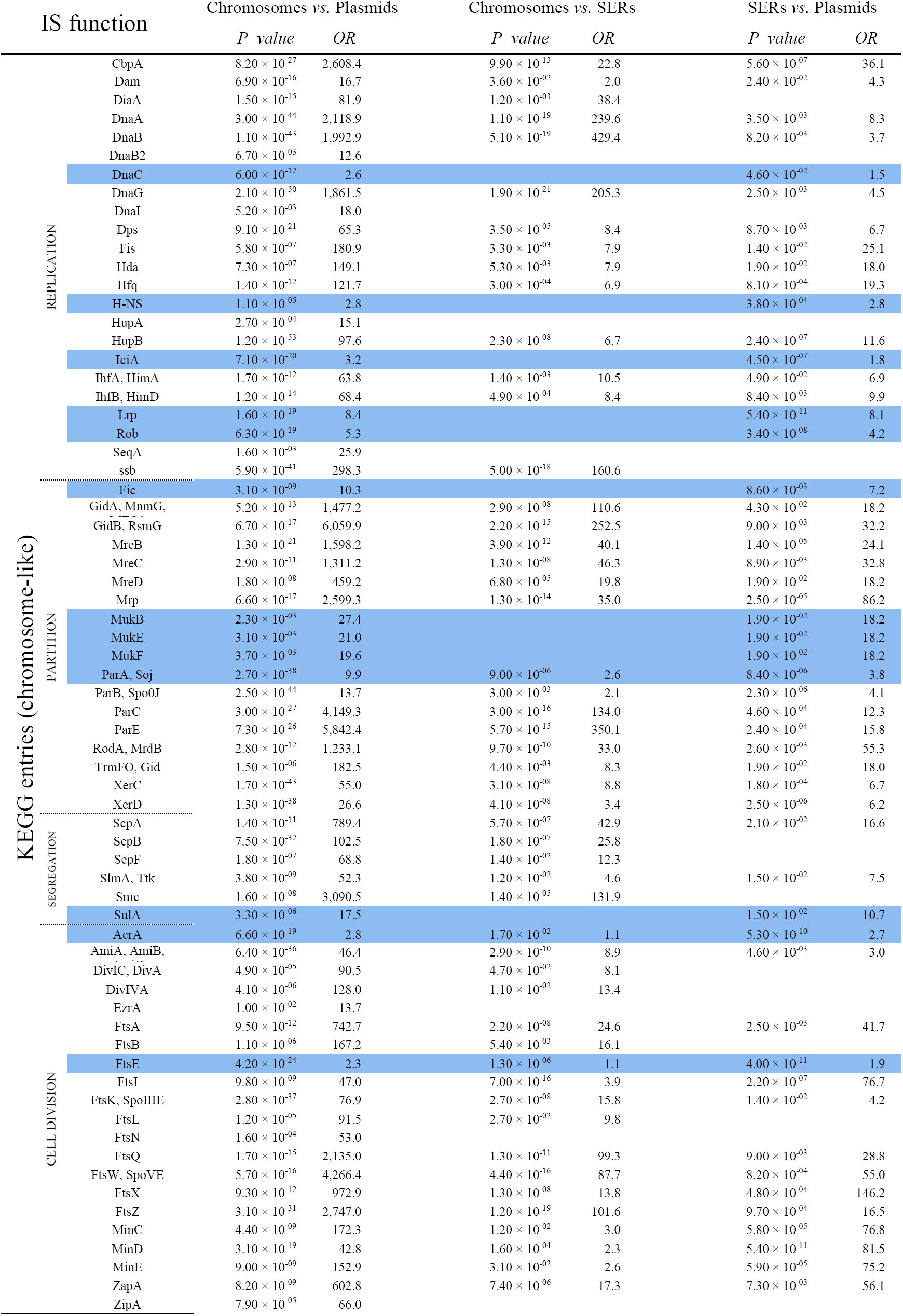
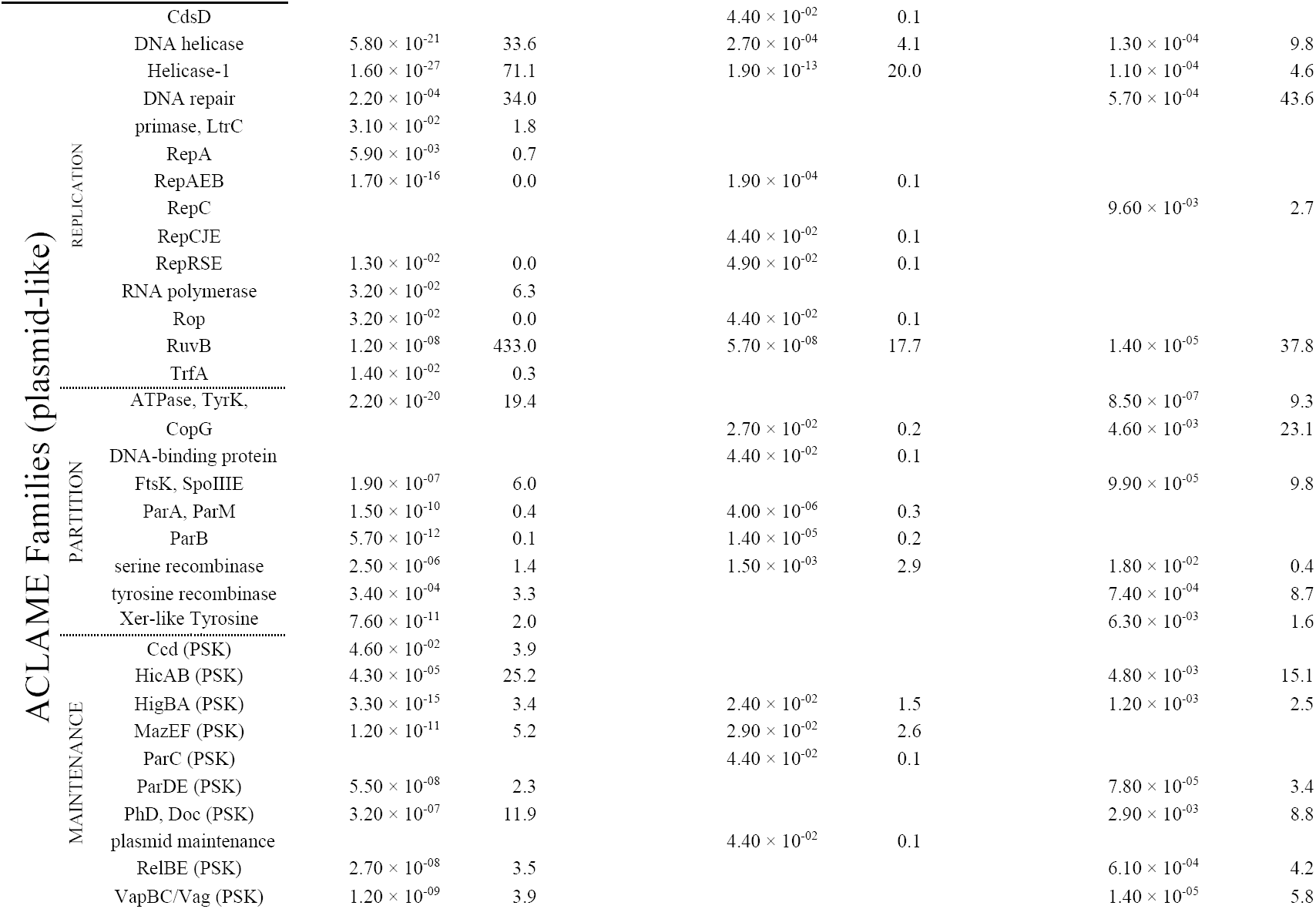
IS usage comparison between replicon categories. Between classes of replicons logistic regressions for each IS function. Model significance: 0 < *P_value* < 0.01: significant; 0.01 < *P_value* < 0.05: poorly significant; 0.05 < *P_value:* non significant (not shown). Odd-ratio (*OR*) favouring the first class: 10^0^ ≤ *OR,* or the second class: *OR* < 10^0^. IS functions biased to the same order of magnitude in chromosomes and SERs when compared to plasmids are highlighted (blue).

Chromosome-signature ISs are also present on the SERs, and some of them are enriched to the same order of magnitude in both classes but not in plasmids (highlighted in Table 6). Among these latter, helicase loader DnaC participates to the replication initiation of the chromosome (Chodavarapu et al., 2016) whilst Walker-type ATPase ParA/Soj interacts with ParB/Spo0J in the *parABS* chromosomal partinioning system, and is required for proper separation of sister origins and synchronous DNA replication (Murray and Errington, 2008). The other ISs have a regulatory role, either locally or globally. Nucleoid-associated proteins (NAPs; Dillon and Dorman, 2010) contribute to the replication regulation: H-NS (histone-like nucleoid structuring protein), IciA (chromosome initiator inhibitor, LysR family transcriptional regulator), MukBEF (condensin), and Rob/ClpB (right arm of the replication origin binding protein/curved DNA-binding protein B, AraC family transcriptional regulator) influence both the conformation and the functions of chromosomal DNA, replication, recombination and repair. The NAPs also have pleiotropic regulatory roles in global regulation of gene transcription depending on cell growth conditions (H-NS, IciA, Lrp (leucine-responsive regulatory protein, Lrp/AsnC family transcriptional regulator), and Rob/ClpB). Similarly, the membrane fusion protein AcrA is a growth-dependent regulator, mostly known for its role as a peripheral scaffold mediating the interaction between AcrB and TolC in the AcrA-AcrB-TolC Resistance-Nodule-cell Division-type efflux pump that extrudes from the cell compounds that are toxic or have a signaling role (Du et al., 2018). It is central to the regulation of cell homeostasis and proper development (Anes et al., 2015; Du et al., 2018; Webber et al., 2009) as well as biofilm formation (Alav et al., 2018). Fic (cell filamentation protein) targets the DNA gyrase B (GyrB) to regulate the cell division and cell morphology (Lu et al., 2018) whereas SulA inhibits FtsZ assembly, hence causing incomplete cell division and filamentation (Chen et al, 2012). FtsE is involved in the Z-ring assembly and the initiation of constriction, and in late stage cell separation (Meier et al, 2017).

The main divergence between SERs and chromosomes lies in the distribution patterns of the ACLAME-annotated ISs (*OR* < 10^0^ in the chromosomes *vs.* SERs comparison). Their higher abundance on the SERs suggests a stronger link of SERs to plasmids. This pattern may also arise from the unbalanced taxon representation in our SER dataset due to a single bacterial lineage. For example, the presence of RepC is likely to be specific to Rhizobiales SERs (Pinto et al., 2012).

### Identification of candidate SERs

Since the IS profiles constitute replicon-type signatures, we searched for new putative SERs or chromosomes among the extra-chromosomal replicons. We used the IS functions as features to perform supervised classification analyses with various training sets (Table 7).

**Table 7.** Performance of the ERT classification procedures.

| TRAINING SET<sup>a</sup> | $CV_{score}$<sup>b</sup> | $\sigma_{CV_{score}}$<sup>c</sup> | $OOB_{score}$<sup>d</sup> | $\sigma_{OOB_{score}}$<sup>e</sup> |
| --- | --- | --- | --- | --- |
| $\{E_{SER}, E'_{plasmid}\}$ | 0.96 | - | 0.96 | - |
| $\{E_{SER}, E'_{plasmid}\}^{it}$ | 0.92 | 0.02 | 0.93 | 0.02 |
| $\{E_{chromosome}, E'_{plasmid}\}$ | 1.00 | - | 1.00 | - |
| $\{E_{SER}, E_{chromosome}\}^{it}$ | 0.98 | 0.00 | 0.98 | 0.01 |

<sup>a</sup> $E_{chromosome}$ and $E_{SER}$ are host genus-normalized sets of chromosomes or SERs, respectively (cf. Table 13). $E'_{plasmid}$ is derived from the INFOMAP clustering solution, by discarding plasmids belonging to clusters also harbouring SERs or chromosomes, and normalized according to host genus. “it” designates the iterative procedure.
<sup>b</sup> Cross-validation score or mean of iteration cross-validation scores.
<sup>c</sup> Standard deviation of iteration cross-validation scores.
<sup>d</sup> Out-of bag estimate or mean of iteration Out-of bag estimates.
<sup>e</sup> Standard deviation of iteration Out-of bag estimates.

The coherence of the SER class (overall high values of the probability for a SER to be assigned to its own class in Tables 7 and 8) confirmed that the ISs are robust genomic markers for replicon characterization. The low SER probability scores presented by a few SERs (Table 8) likely result from a low number of carried ISs (*e.g., Leptospira*), or from the absence in the data of lineage-specific ISs (e.g., SER idiosyncratic replication initiator RtcB of Vibrionaceae).

**Table 8.** SER probability to belong to the *SER* class.

| Genus | $\bar{P}_{SER}(SER)^a$ |
| --- | --- |
| *Agrobacterium* | 0.90 |
| *Aliivibrio* | 0.87 |
| *Anabaena* | 0.94 |
| *Asticcacaulis* | 0.95 |
| *Brucella* | 0.92 |
| *Burkholderia* | 0.89 |
| *Butyrivibrio* | 0.83 |
| *Chloracidobacterium* | 0.88 |
| *Cupriavidus* | 0.94 |
| *Cyanothece* | 0.86 |
| *Deinococcus* | 0.78 |
| *Ensifer/Sinorhizobium* | 0.90 |
| *Ilyobacter* | 0.88 |
| *Leptospira* | 0.54 |
| *Nocardiopsis* | 0.90 |
| *Ochrobactrum* | 0.96 |
| *Paracoccus* | 0.96 |
| *Photobacterium* | 0.95 |
| *Prevotella* | 0.92 |
| *Pseudoalteromonas* | 0.91 |
| *Ralstonia* | 0.95 |
| *Rhodobacter* | 0.69 |
| *Sphaerobacter* | 0.88 |
| *Sphingobium* | 0.73 |
| *Thermobaculum* | 0.78 |
| *Variovorax* | 0.83 |
| *Vibrio* | 0.76 |

<sup>a</sup> SER probability, averaged per host genus, to belong to the *SER* class in the supervised classification using $\{E_{SER}, E_{plasmid}\}^{it}$ as training set.

We detected a number of candidate SERs among the plasmids (Table 9a), most of which are essential to the cell functioning and/or the fitness of the organism (*cf.* Box 1). Whereas most belong to bacterial lineages known to harbour multipartite genomes, novel taxa emerge as putative hosts to complex genomes (Rhodospirillales and, to a lesser extent, Actinomycetales). In contrast, our analyses confirmed only one putative SER (*Ruegeria* sp. TM1040) within the *Roseobacter* clade (Petersen et al., 2013). Remarkably, we identified eight candidate chromosomes corresponding to two plasmids, also identified as candidate SERs, that encode ISs hardly found in extra-chromosomal elements (*e.g.,* DnaG, DnaB, ParC and ParE), and six SERs that part of, or all, our analyses associate to standard chromosomes (Table 9b). Notably, *Prevotella intermedia* SER (CP003503) shows a very high probability (> 0.98) to be a chromosome while its annotated chromosome (CP003502), unique of its kind, falls within the plasmid class. This approach can thus be extended to test the type of replicon for (re)annotation purposes.

**Table 9.**
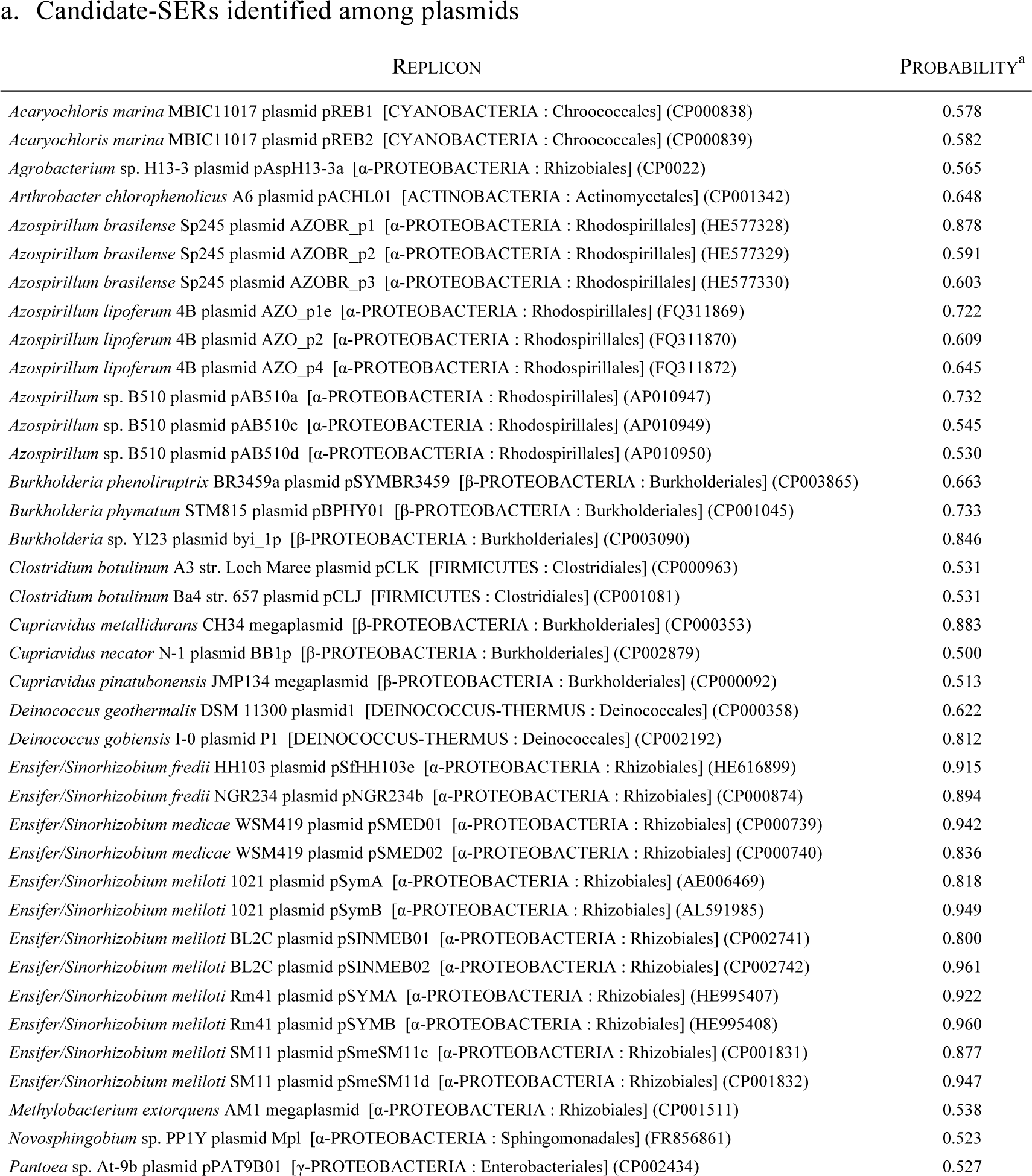
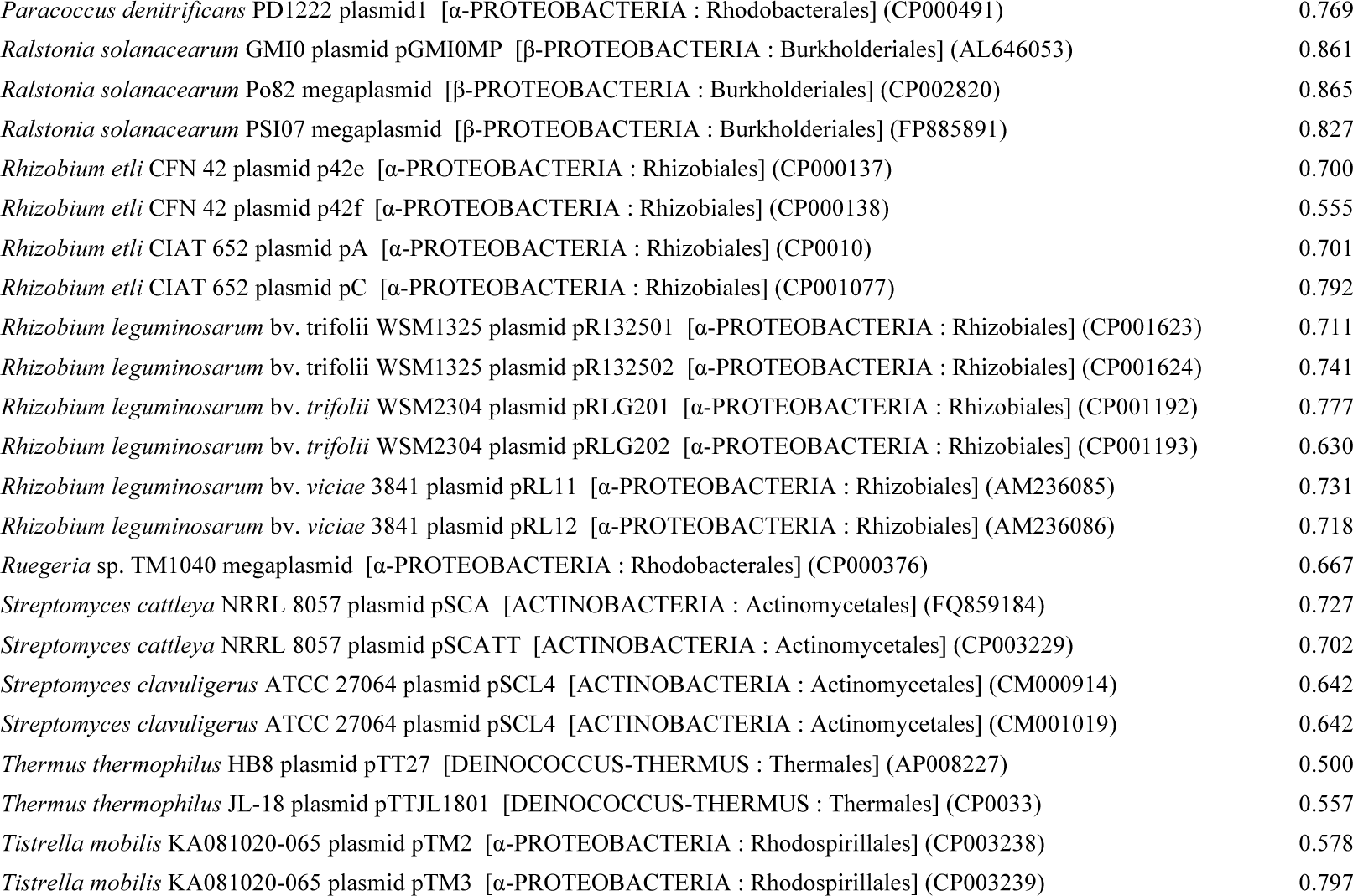
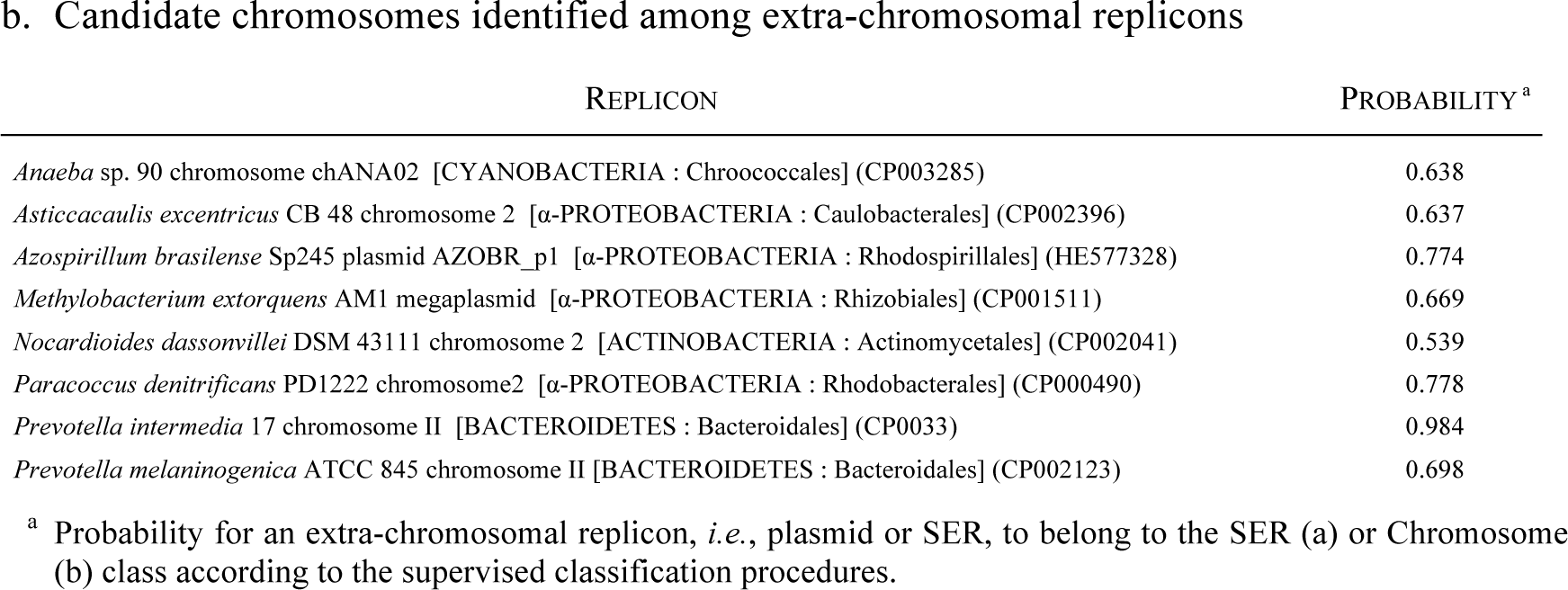
Identification of ERs among the extra-chromosomal replicons.

##### BOX 1. CHARACTERISTICS OF CANDIDATE SERS

According to the literature, most candidate SERs that we detected among plasmids (Table 9a) were expected to be essential to the cell functioning and/or to the fitness of the organism.

- *Azospirillum* genomes are constituted of multiple replicons, at least one of which is expected to be essential. The largest extra-chromosomal replicon in *A. brasilense* was proposed to be essential for bacterial life (Wisniewski-Dyé et al., 2011) since it encodes well-conserved housekeeping genes involved in DNA replication, RNA metabolism and biosynthesis of nucleotides and cofactors, as well as in transport and protein post-translational modifications. This replicon is unambiguously identified as a SER by our analyses, as are additional replicons found in *A. lipoferum* and *A.* sp. B510, expected homologues to *A. brasilense* SER (Acosta-Cruz et al., 2012). In contrast, other extra-chromosomal replicons classified as chromids by Wisniewski-Dyé et al. (2012) are unlikely to be true essential replicons. They were not retrieved among our candidate SERs.
- In *Rhizobium etli* CFN42, functional interactions among sequences scattered in the different extrachromosomal replicons are required for successful completion of life in symbiotic association with plant roots or saprophytic growth (Brom et al., 2000). p42e (CP000137) is the only replicon other than the chromosome that contains genes involved in the primary metabolism (Landeta et al., 2011; Villaseñor *et al.* 2011) and evades its elimination by co-integration with other replicons including the chromosome (Landeta et al., 2011). Furthermore, homologues to this replicon were identified in the genomes of other *R. etli* strains as well as other *Rhizobium* species: *R. etli* CIAT652 pA, *R. leguminosarum* bv. *viciae* 3841 pRL11, *R. leguminosarum* bv. *trifolii* WSM2304 pRLG202 and *R. leguminosarum* bv. *trifolii* WSM1325 pR132502 (CP001075, AM236085, CP001193, and CP001624, respectively) (Landeta et al., 2011; Villaseñor et al., 2011). These replicons were thus proposed to be secondary chromosomes (Landeta et al., 2011).
- The genome of *Ensifer/Sinorhizobium meliloti* AK83 was the single multipartite-annotated *Ensifer/Sinorhizobium* genomes present in our dataset. This bacterium carries two large extra-chromosomal replicons that are involved in the establishment of the nitrogen fixation symbiosis with legume plants. pSymA contains most of the genes involved in the nodulation and nitrogen fixation whereas pSymB carries exopolysaccharide biosynthetic genes, also required for the establishment of the symbiosis. Our analyses identifies candidate SERs similar to *S. meliloti* AK83 pSymA and pSymB in other *S. meliloti* strains as well as in *S. fredii* and *S. medicae.* pSymB has been referred to as second chromosome for carrying genes encoding essential house-keeping functions (Blanca-Ordóñez et al., 2010; Galardini et al., 2011). It shows a higher level of conservation across strains and species than pSymA (Galardini et al., 2013). pSymA, generally thought to be as stable as pSymB, greatly contribute to the bacterial fitness in the rhizosphere (Blanca-Ordóñez et al., 2010; Galardini et al., 2013).
- The identification of *Methylobacterium extorquens* AM1 1.2 Mb megaplasmid as a SER is supported by its presence in the genome in a predicted one copy number, by its coding a truncated *luxI* gene essential for the operation of two chromosomally-located *luxI* genes, as well as the single *umuDC* cluster involved in SOS DNA repair, and by the presence of a 130 kb region syntenic to a region of similar length in the chromosome of *Methylobacterium extorquens* strain DM4 (Vuilleumier et al., 2009).
- The megaplasmid (821 kb) in *Ruegeria* sp. TM1040 carries more rRNA operons (3) than the chromosome (1) and several unique genes (Moran et al., 2007). *Ruegeria* sp. TM1040 is the only species in the *Roseobacter* group that possesses a SER. None of the plasmids in the other species included in our datasets qualified as SERs according to our results in contrast to the commonly held view (Petersen et al., 2013).
- In *Burkholderia* genus, additional ERs possess a centromere whose sequence is distinct from, but strongly resembles that of the chromosome centromere (Dubarry et al., 2009). However, these centromeres have a common origin and a plasmid ancestry (Passot et al., 2012). The evolution of these replicons into SERs is best accounted for by the high level of plasticity observed in the *Burkholderia* genomes, with extra-chromosomal replicons going through extensive exchange of genetic material among them as well as with the chromosomes (Maida et al., 2014).
- *Acaryochloris marina* MBIC11017 pREB1 (CP000838) and pREB2 (CP000839) plasmids were identified as candidate SERs. Both these megaplasmids code for metabolic key-proteins, and are thus likely to contribute to the bacterium fitness (Swingley et al., 2008).
- The genomes of *Streptomyces cattleya* NRRL8057 and *S. clavuligerus* ATCC27064 harbour a linear megaplasmid (1.8 Mb) that shows a high probability (P ≈ 0.7) to be a SER. The megaplasmid of *S. cattleya* NRRL8057 encodes genes involved in the synthesis of various antibiotics and secondary metabolites and is expected to be important to the life of the bacterium in its usual habitat (Barbe et al., 2011; O’Rourke et al., 2009). In *S. clavuligerus* ATCC27064, none of the megaplasmid-encoded genes are expected to belong to the core genome (Medema et al., 2010). However, the megaplasmid is likely to contribute to the bacterium firness. It represents more than 20% of the coding genome and constitutes a large reservoir of genes involved in bioactive compound production and cross-regulation with the chromosome (Medema et al., 2010). Furthermore, *S. clavuligerus* chromosome requires the SER-encoded *tap* gene involved in the telomere replication.
- *Butyrivibiro proteoclasticus* B316 harbours two plasmid, one of which, pCY186 plasmid (CP001813), was identified as a candidate SERs by our analysis, albeit with a low probability (0.56). In support to this, it carries numerous genes coding for proteins involved in replication of the chromosome (Yeoman et al., 2011). The second plasmid in that strain, pCY360 (CP001812), also proposed to be an essential replicon in that bacterium (Yeoman et al., 2011), presents too low a probability (P = 0.32) in our analysis to qualify as a SER.

## DISCUSSION

The SERs clearly stand apart from plasmids, including those that occur consistently in a bacterial species, *e.g., Lactobacillus salivarius* pMP118-like plasmids (Li et al., 2007). The replicon size proposed as a primary classification criterion to separate the SERs from the plasmids (diCenzo and Finan, 2017; Harrison et al., 2010) proves to be inoperative. The IS profiles accurately identify the SERs of *Leptospira* and *Butyrivibrio* despite their plasmid-like size, and unambiguously ascribe the chromosomes in the reduced genomes of endosymbionts (sizes down to 139 kb) to the chromosome class. Conversely, they assign *Rhodococcus jostii* RHA1 1.12 Mb-long pRHL1 replicon to the plasmid class, and do not discriminate the megaplasmids (>350 kb (diCenzo and Finan, 2017)) from smaller plasmids. Plasmids may be stabilized in a bacterial population by rapid compensatory adaptation that alleviates the fitness cost incurred by their presence in the cell (San Millan et al., 2014; Hall et al., 2017; Stalder et al., 2017). This phenomenon involves mutations either on the chromosome only, on the plasmid only, or both, and does not preclude the segregational loss of the plasmid. On the contrary, SERs code for chromosome-type IS proteins that integrate them constitutively in the species genome and the cell cycle. The SERs thence qualify as essential replicons regardless of their size and of the phenotypical/ecological, possibly essential, functions that they encode and which vary across host taxa.

Yet, SERs also carry plasmid-like ISs, suggesting a role for plasmids in their formation. The prevailing opinion assumes that SERs derive from the amelioration of megaplasmids (diCenzo and Finan, 2017; diCenzo et al., 2013; Harrison et al., 2010; MacLellan et al., 2004; Slater et al., 2009): a plasmid bringing novel functions for the adaptation of its host to a new environment is stabilized into the bacterial species genome through the transfer from the chromosome of essential genes (diCenzo and Finan, 2017; Slater et al., 2009). However, the generalized presence of chromosome-like ISs in the SERs of the various taxa with multipartite genomes is unlikely to derive from the action of environment-specific and lineage-specific selective forces. In reverse, all bacteria with similar lifestyle and exhibiting some phylogenetic relatedness may not harbor multiple ERs (e.g., α-proteobacterial nitrogen-fixing legume symbionts). Also, the gene shuttling from chromosome to plasmid proposition fails to account for the situation met in the multipartite genomes of *Asticaccaulis excentricus, Paracoccus denitrificans* and *Prevotella* species. Their chromosome-type ISs are evenly distributed between the chromosome and the SER whereas their homologues in the mono- or multipartite genomes of most closely related species are primarily chromosome-coded (see Table 10 for an example). This pattern, mirrored in their whole gene content (Naito et al., 2016; Poirion, 2014), hints at the stemming of the two essential replicons from a single chromosome by either a splitting event or a duplication followed by massive gene loss. Neither mechanism informs on the presence of plasmid-type maintenance machinery on one of the replicons. The severing of a chromosome generates a single true replicon carrying the chromosome replication origin and an origin-less remnant, whilst the duplication of the chromosome produces two chromosomal replicons with identical maintenance systems. Whereas multiple copies of the chromosome are known to cohabit constitutively in polyploid bacteria (Ohtani et al., 2010), the co-occurrence of dissimilar chromosomes bearing identical replication initiation and partition systems is yet to be described in bacteria.

**Table 10.** IS profiles of *Paracoccus denitrificans vs. Rhodobacter sphaeroides* (Rhodobacterales) Chromosome-like IS functions coded only by the SER in *P. denitrificans* or *R. sphaeroides* whilst by the chromosome in sther Rhodobacterales are indicated by an asterisk. Numbers corresponds to the number of homologues (*P. denitrificans* PD1222) or the pourcentage of function-coding replicons (*R. sphaeroides* and Rhodobacterales genomes).

|  |  | *Paracoccus denitrificans* PD1222 |  |  | *Rhodobacter sphaeroides* |  |  | Other Rhodobacterales |  |
| --- | --- | --- | --- | --- | --- | --- | --- | --- | --- |
|  |  | Chromosome1 (CP000489) | Chromosome2 (CP000490) | plasmid1 (CP000491) | Chromosome % (n=12) | SER % (n=12) | plasmid % (n=14) | Chromosome % (n=12) | plasmid % (n=33) |
| KEGG entry (chromosome-like) | IS FUNCTION GROUP |  |  |  |  |  |  |  |  |
|  | CbpA | 1 |  |  | 100 |  |  | 75 |  |
|  | Dam | 1 |  |  | 50 | 75 |  | 25 |  |
|  | DnaA | * | 1 |  | 100 |  |  | 92 |  |
|  | DnaB | * | 1 |  | 100 |  |  | 100 |  |
|  | DnaC |  |  |  | 25 |  |  |  | 3 |
|  | DnaG | * | 1 |  | 100 |  |  | 100 |  |
|  | Dps | 1 |  |  |  |  |  | 50 | 15 |
|  | Fis | * | 1 |  | 100 |  |  | 75 |  |
|  | Hda |  |  |  |  |  |  | 42 |  |
|  | Hfq | * | 1 |  | 100 |  |  | 100 |  |
|  | H-NS | 2 | 2 |  | 100 |  | 29 | 67 | 9 |
|  | HupA |  |  |  |  |  |  | 17 |  |
|  | HupB | 1 |  |  | 100 |  |  | 100 |  |
|  | IciA | 1 |  |  |  |  | 36 | 67 |  |
|  | IhfA, HimA | 1 |  |  | 100 |  |  | 100 |  |
|  | IhfB, HimD | * | 1 |  | * | 100 |  | 100 |  |
|  | Lrp | 4 | 1 | 3 | 100 | 25 | 50 | 100 | 9 |
|  | Rob |  |  |  | 25 |  |  | 8 |  |
|  | ssb | 1 | 3 |  | 100 | 100 |  | 100 | 3 |
|  | Fic |  |  |  |  |  | 7 | 8 | 6 |
|  | GidA, MnmG, MTO1 | 1 |  |  | 100 |  |  | 100 |  |
|  | GidB, RsmG | 1 |  |  | 100 |  |  | 100 |  |
|  | MreB | * | 1 |  | 100 |  |  | 92 |  |
|  | MreC | * | 1 |  | 100 |  |  | 92 |  |
|  | MreD | * | 1 |  | 100 |  |  | 92 |  |
|  | Mrp | 1 |  |  | 100 |  |  | 100 |  |
|  | ParA, Soj | 4 | 1 |  | 100 |  |  | 100 | 9 |
|  | ParB, Spo0J | 3 | 1 | 1 | 100 | 75 |  | 100 | 42 |
|  | ParC | 1 | 1 |  | 100 |  |  | 100 |  |
|  | ParE | 1 |  |  | 100 |  |  | 100 |  |
|  | RodA, MrdB | * | 1 |  | 100 |  |  | 92 |  |
|  | TrmFO, Gid | * | 1 |  | 100 |  |  | 100 |  |
|  | XerC | 1 |  |  | 100 |  |  | 100 |  |
|  | XerD | 1 |  |  | 100 | 25 | 7 | 100 | 6 |
|  | ScpA | 1 |  |  | 100 |  |  | 100 |  |
|  | ScpB | 1 |  |  | 100 |  |  | 100 |  |
|  | Smc | 1 |  |  | 100 |  |  | 100 |  |
|  | AcrA | 2 | 2 |  | * | 100 | 7 | 58 |  |
|  | AmiA, AmiB, AmiC | 1 |  |  | 100 |  |  | 100 |  |
|  | FtsA | * | 1 |  | 100 |  |  | 83 |  |
|  | FtsE | 1 |  |  | 100 |  | 7 | 92 |  |
|  | FtsI | 1 |  |  | 100 |  |  | 92 |  |
|  | FtsK, SpoIIIE | 1 |  |  | 100 |  |  | 100 |  |
|  | FtsQ | * | 1 |  | 100 |  |  | 92 |  |
|  | FtsW, SpoVE |  | 1 |  | 100 |  |  | 100 |  |
|  | FtsX | 1 |  |  | 100 |  | 7 | 92 |  |
|  | FtsZ | * | 1 |  | 100 |  |  | 100 |  |
|  | MinC |  |  | 1 |  |  |  |  |  |
|  | MinD |  |  | 1 | 75 |  |  | 75 |  |
|  | MinE |  |  | 1 |  |  |  |  |  |
|  | ZapA | 1 |  |  | 100 |  |  | 100 |  |
| ACLAME family (plasmid-like) | IS FUNCTION GROUP |  |  |  |  |  |  |  |  |
|  | DNA helicase |  |  |  | 25 | 50 | 7 | 25 |  |
|  | Helicase-1 | 1 | 1 |  | 100 | 50 |  | 92 | 3 |
|  | Helicase-2 |  |  |  |  |  |  | 8 | 3 |
|  | DNA repair |  |  |  |  |  |  | 8 |  |
|  | RepA | 2 | 2 |  |  |  | 43 |  | 42 |
|  | RepAEB |  |  | 1 |  | 100 | 7 |  | 15 |
|  | RepC | 1 |  |  | 50 |  | 43 | 8 | 24 |
|  | RuvB | 1 |  |  | 100 |  | 7 | 92 |  |
|  | ATPase, TyrK, ExoP | 2 |  |  | 100 | 50 | 14 | 75 | 15 |
|  | ParA, ParM |  | 1 | 1 |  | 100 | 100 | 8 | 88 |
|  | ParB |  |  |  | 75 | 100 | 93 | 25 | 64 |
|  | plasmid dimer resolution |  |  |  |  |  | 14 |  | 15 |
|  | Serine recombinase | 3 |  |  | 25 |  | 50 | 50 | 30 |
|  | Tyrosine recombinase |  |  |  | 25 |  | 7 | 33 | 6 |
|  | Xer-like tyrosine recombinase |  | 1 |  | 25 | 50 |  | 33 | 6 |
|  | XerD |  |  |  | 25 |  |  |  |  |
|  | Ccd (PSK) |  |  |  |  |  | 7 |  |  |
|  | HicAB (PSK) |  |  |  |  |  | 7 |  |  |
|  | HigBA (PSK) | 2 |  |  |  |  | 14 | 17 | 6 |
|  | MazEF (PSK) |  |  |  |  |  | 7 |  | 6 |
|  | ParDE (PSK) | 2 | 6 |  | 25 | 50 | 7 | 8 | 15 |
|  | PhD, Doc (PSK) |  |  |  |  |  | 7 |  |  |
|  | RelBE (PSK) | 1 |  |  |  |  |  |  | 3 |
|  | VapBC/Vag (PSK) | 3 | 2 |  |  |  | 14 | 8 | 9 |

We propose that the requirement for maintenance system compatibility between co-occurring replicons is the driving force behind the presence of plasmid-type replication initiation and maintenance systems in bacterial SERs. Indeed, genes encoding chromosome-like replication initiators (DnaA) are hardly found on SERs. When they are, in *Paracoccus denitrificans, Prevotella intermedia* and *P. melaninogenica,* the annotated chromosome in the corresponding genome does not carry one. Similarly, chromosomal centromeres (*parS*) are found on a single replicon within a multipartite genome, which is the chromosome in all genomes but one. In *P. intermedia* (GCA_000261025.1), both replication initiation and partition systems define the SER as the *bona fide* chromosome and the annotated chromosome as an extra-chromosomal replicon. The harmonious coexistence of different replicons in a cell requires that they use divergent enough maintenance systems. In the advent of a chromosome fission or duplication, the involvement of an autonomously self-replicating element different from the chromosome is mandatory to provide one of the generated DNA molecules with a (non-chromosomal) maintenance machinery.

‘Plasmid-first’ and ‘chromosome-first’ hypotheses can be reconciled into a unified, general Fusion-Shuffling-Scission model of SER emergence where a chromosome and a plasmid combine into a cointegrate (Fig. 6). Plasmids are known to merge or to integrate chromosomes in both experimental settings (Brom et al., 2000; Guo et al., 2003; Iordânescu, 1975; Sýkora, 1992) and the natural environment (Cervantes et al., 2011; Naito et al., 2016; Sýkora, 1992), as are the SER and chromosome of a multipartite genome (Val et al., 2014; Xie et al., 2017; Yamamoto et al., 2018). When integrated, the plasmids/SERs can thus replicate with the chromosome and persist in the bacterial lineage through several generations (Cervantes et al., 2011; Val et al., 2014; Xie et al., 2017). The co-integrate may resolve into its original components (Guo et al., 2003; Val et al., 2014) or give rise to novel genomic architectures (Guo et al., 2003; Cervantes et al., 2011; Val et al., 2014). The co-integration state likely facilitates inter-replicon gene exchanges and genome rearrangements that may lead to the translocation of large chromosome fragments to the resolved plasmid (Guo et al., 2003; Val et al., 2014). Multiple cell divisions, and possibly several merging-resolution rounds, could provide time and opportunity for the plasmid-chromosome re-assortment to take place, and for multiple essential replicons and a viable distributed genome to form ultimately. In the novel genome, one ER retains the chromosome-like origin of replication and centrosome, and the other the plasmidic counterparts. The novel ERs differ from the chromosome and plasmid that gathered in the progenitor host at the onset. They thus constitute neo-chromosomes that carry divergent maintenance machineries and can cohabit and function in the same cell. Depending on the number of cell cycles spent as co-integrate, the level of genome reorganization, the acquisition of genetic material and the environmental selective pressure acting upon the host, the final essential replicons may exhibit diverse modalities of genome intégration (Figure 6).

**Figure 6.**
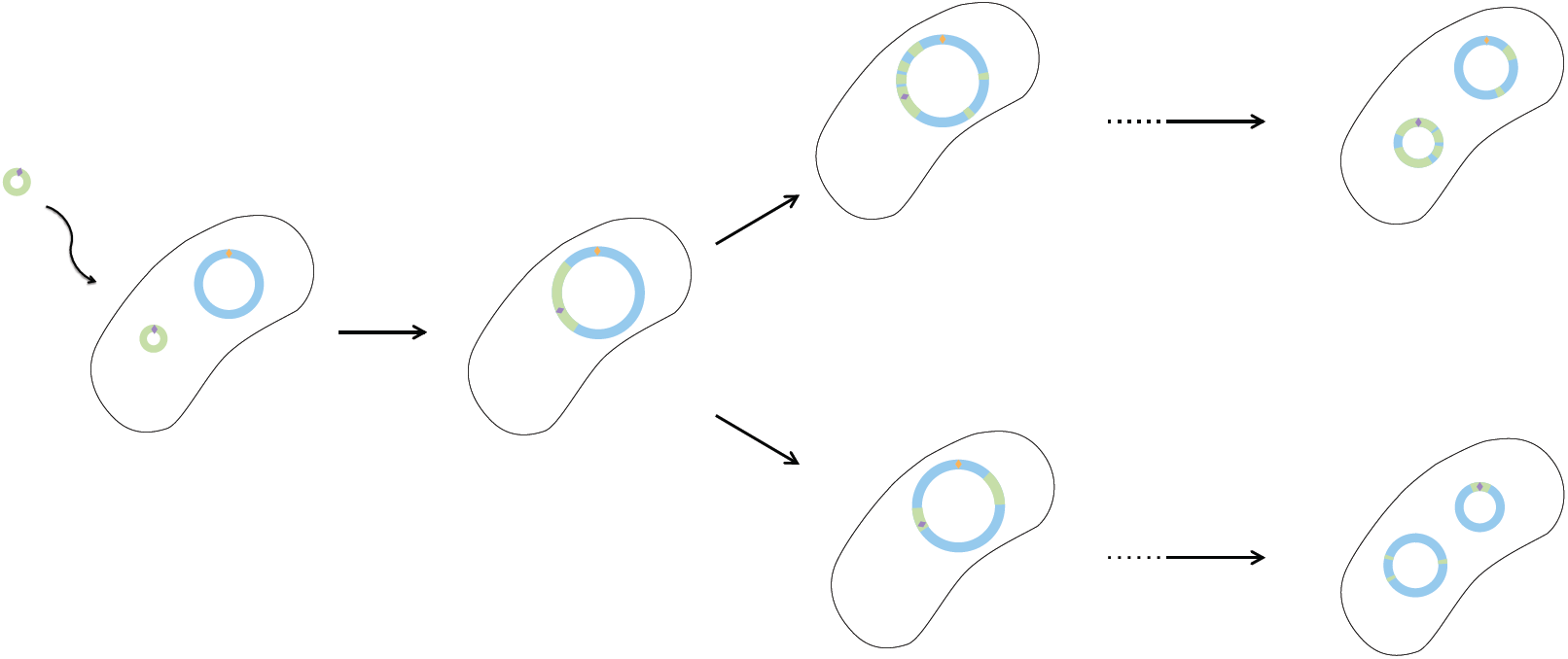
Fusion-Shuffling-Scission model of distributed genome evolution. Origins of replication are represented by diamonds.

The Fusion-Shuffling-Scission model of genome evolution that we propose accounts for the extreme plasticity met in distributed genomes and the eco-phenotypic flexibility of their hosts. Indeed, having a distributed genome appears to extend and accelerate the exploration of the genome evolutionary landscape, producing complex regulation (diCenzo et al., 2018; Galardini et al., 2015; Jiao et al., 2018) and leading to novel eco-phenotypes and species diversification (e.g., Burkholderiaceae and Vibrionaceae). Furthermore, this model may explain the observed separation of the replicons according to taxonomy. Chromosomes and plasmids thus appear as extremes on a continuum of a lineage-specific genetic material.

## MATERIALS AND METHODS

To understand the relationships between the chromosomal and plasmidic replicons, we focused on the distribution of Inheritance System (IS) genes for each replicon and built networks linking the replicons given their IS functional orthologues (Fig. 2).

### Retrieval of IS functional homologues

A sample of proteins involved in the replication and segregation of bacterial replicons and of the bacterial cell cycle was constructed using datasets available from the ACLAME (Leplae et al., 2010) and KEGG (Kanehisa et al., 2012) databases. Gene ontologies for “replication”, “partition”, “dimer resolution”, and “genome maintenance” (Table 11) were used to select related ACLAME plasmid protein families (Table 1) using a semi-automated procedure.

**Table 11.**
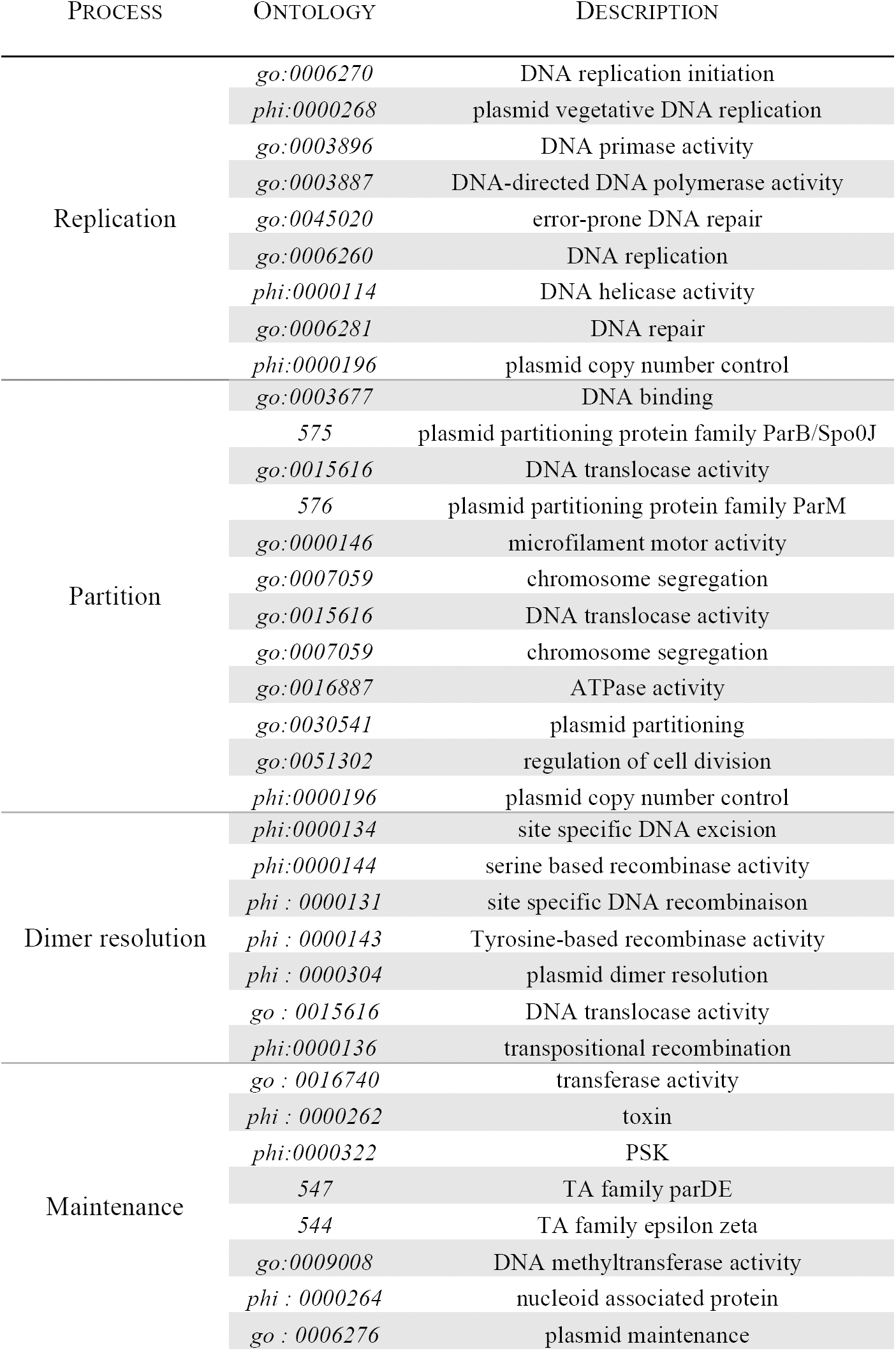
Gene ontologies related to plasmid ISs used to select groups of orthologous proteins from the ACLAME database

KEGG orthology groups were selected following the KEGG BRITE hierarchical classification (Table 2). Then, the proteins belonging to the relevant 92 ACLAME protein families and 71 KEGG orthology groups (3,847 and 43,757 proteins, respectively) were retrieved and pooled. Using this query set amounting to a total of 47,604 proteins, we performed a *blastp* search of the 6,903,452 protein sequences available from the 5,125 complete sequences of bacterial replicons downloaded from NCBI Reference Sequence database (RefSeq) (Pruitt et al., 2007) on 30/11/2012. We identified 358,624 putative homologues using BLAST default parameters (Camacho et al., 2009) and a 10^-5^ significance cut-off value. We chose this *E*-value threshold to enable the capture of similarities between chromosome and plasmid proteins whilst minimizing the production of false positives, *i.e.,* proteins in a given cluster exhibiting small *E*-values despite not being functionally homologous. Using RefSeq ensured the annotation consistency of the genomes included in our dataset.

### Clustering of Is functional homologues

Using this dataset, we inferred clusters of IS functional homologues by coupling of an *all-versus-all blastp* search using a 10^-2^ *E*-value threshold and a TRIBE-MCL (Enright et al., 2002) clustering procedure. As input to TRIBE-MCL, we used the matrix of log transformed *E*-value, *d*(*p*_*j*_, *p*_*j*_) = - log_10_ (*e*_*value*_(*p*_*i*_, *p*_*j*_)), obtained from the comparisons of all possible protein pairs. Using a granularity value of 4.0 (see below), we organized the 358,624 IS homologues into 7013 clusters, each comprising from a single to 1990 proteins (Figure 3). We annotated IS homologues according to their best match (BLAST hit with the lowest *E*-value) among the proteins of the query set, *i.e.,* according to one of the 117 functions of the query set (71 from KEGG and 46 from ACLAME). Then, we named the clusters of functional homologues using the most frequent annotation among the proteins in the cluster. We used the number of protein annotations in a cluster to determine the cluster quality, a single annotation being optimal. To select the best granularity and to estimate the consistency of the clusters in terms of functional homologues, we computed the weighted Biological Homogeneity Index (*wBHI,* modified from the *BHI* (Datta and Datta, 2006), each cluster being weighted by its size) and the Conservation Consistency Measure (*CCM*, similar to the *BHI* but using the functional domains of the proteins to define the reference classes), which both take into account the size distribution of the clusters (See next paragraph for details on index calculation). The former gives an estimation of the overall consistency of clusters annotations according to the protein annotations whereas the latter gives an estimation of cluster homogeneity according to the protein domains identified beforehand. To build the sets of functional domains, we performed an *hmmscan* (Finn et al., 2011) procedure against the Pfam database (Finn et al., 2016) of each of the 358,624 putative IS homologues. We annotated each protein according to the domain match(es) with *E*-value < 10^-5^ (individual *E*-value of the domain) and *c-E*-value < 10^-5^ (conditional *E-* value that measures the statistical significance of each domain). If two domains overlapped, we only considered the domain exhibiting the smallest *E*-value. We estimated *wBHI* and *CCM* indices for the clustering of the IS homologues and compared with values obtained for random clusters simulated according to the cluster size distribution of the IS proteins, irrespective of their length or function. For each of the clustering obtained for different granularities, we constructed a random clustering following the original cluster size distribution (assessed with a χ^2^ test) and composed with simulated proteins according to the distributions of the type and number of functional domains of the data collected from the 358,624 IS homologues. Overall, the clusters obtained using a granularity of 4.0 with the TRIBE-MCL algorithm appeared to be homogenous in terms of proteins similarities toward their best BLAST hits and their functional domain distributions (see below).

### Evaluation of the clustering procedures

In order to select the best granularity and to estimate the consistency of the clusters in terms of functional homologs, we computed the *weighted Biological Homogeneity Index* (*wBHI*) and the *Conservation Consistency Measure* (*CCM*). The former gives an estimate of the overall consistency of clusters annotations according to the protein annotations whereas the latter gives an estimate of cluster homogeneity according to protein domains identified beforehand. Although close to the *Biological Homogeneity Index* (*BHI*) introduced by Datta and Datta (2006), both these indices take into account the size distribution of the clusters.

The *BHI* was originally introduced to measure the biological homogeneity of clusters according to reference classes to evaluate clusters obtained with microarray data (Datta and Datta, 2006). Given a clustering *C=*{*C*_*1*_*,…,C*_*k*_} of *k* clusters with *n_i_* the size of the cluster *C_i_*, a set of *m* proteins *P=*{*P*_*1*_*,…,P*_*m*_} and a set *r* of reference classes *R* where each class *R*_*i*_ could be linked to the *m* proteins, the BHI is defined as:

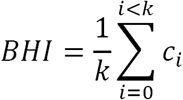

where *c*_*i*_ is defined as:

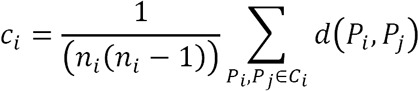

where *d*(*P*_*i*_*,P*_*j*_)*=1* if *P*_*i*_ and *P*_*j*_ share at least one common reference class, and *d*(*P*_*i*_*,P*_*j*_)=0 otherwise. The reference classes here are the annotations defined according to the protein best BLAST hit. The *BHI* is thus an easy-to-interpret measure, which value is maximal when, for all clusters, all the proteins in a cluster share at least one annotation. The *wBHI* is a modification of the *BHI*, where each cluster is weighted by its size *m*. Following the previous notation scheme, the *wBHI* is defined as:

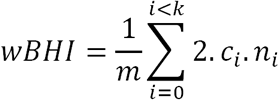

The *CCM* is similar to the *BHI* but the functional domains of the proteins are used to define the reference classes. The distance between the proteins is here computed as the Jaccard distance between the functional domain sets of the proteins. Every protein *P*_*i*_ can be described as a vector of functional domains, *D*_*Pi*_ *=*{*d*_*1*_*, …, d_x_*}. The Jaccard distance between the two sets of domains *d*_*2*_ (*P*_*i*_*P*_*j*_) can be defined as:

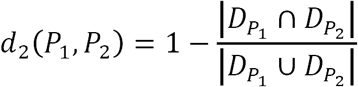

where 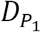 and 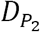 are the clans or domains (when no clan could be assigned) identified in *P* _1_ and *P*_2_ respectively. For a given cluster *C*_*i*_, the *CCM*is calculated as:

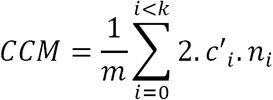

where *c*’_*i*_ is defined as:

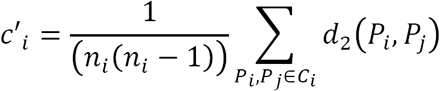

Clusters which proteins have similar domains result in a *CCM* value close to 0, whereas a *CCM* value close to 1 indicates that the clusters hold proteins with little domain overlap.

### Choice of the clustering granularity

We tested several levels of granularity to optimize the TRIBE-MCL clustering and obtain the most informative IS clustering in terms of functional linkage. Too low a granularity would produce large clusters containing multiple functional families. In turn, increasing the granularity results in the tightening of the cluster. A high granularity tends to split clusters harboring different protein subfamilies (*e.g.,* a cluster composed of proteins from the tyrosine recombinase superfamily) and to produce multiple clusters of proteins belonging to a single function family according to their level of sequence dissimilarity. Furthermore, too high a granularity would result in the formation of numerous single protein clusters, and would dramatically increase the computation times of the following analyses. A granularity level of 4.0 constituted a good compromise (Figure 8). Values of *CCM* and *BHI* are slightly improved compared to granularities of 2.0 and 3.0, and the high but still workable number of clusters is expected to prevent the formation of clusters mingling distinct protein subfamilies.

**Figure 8.**
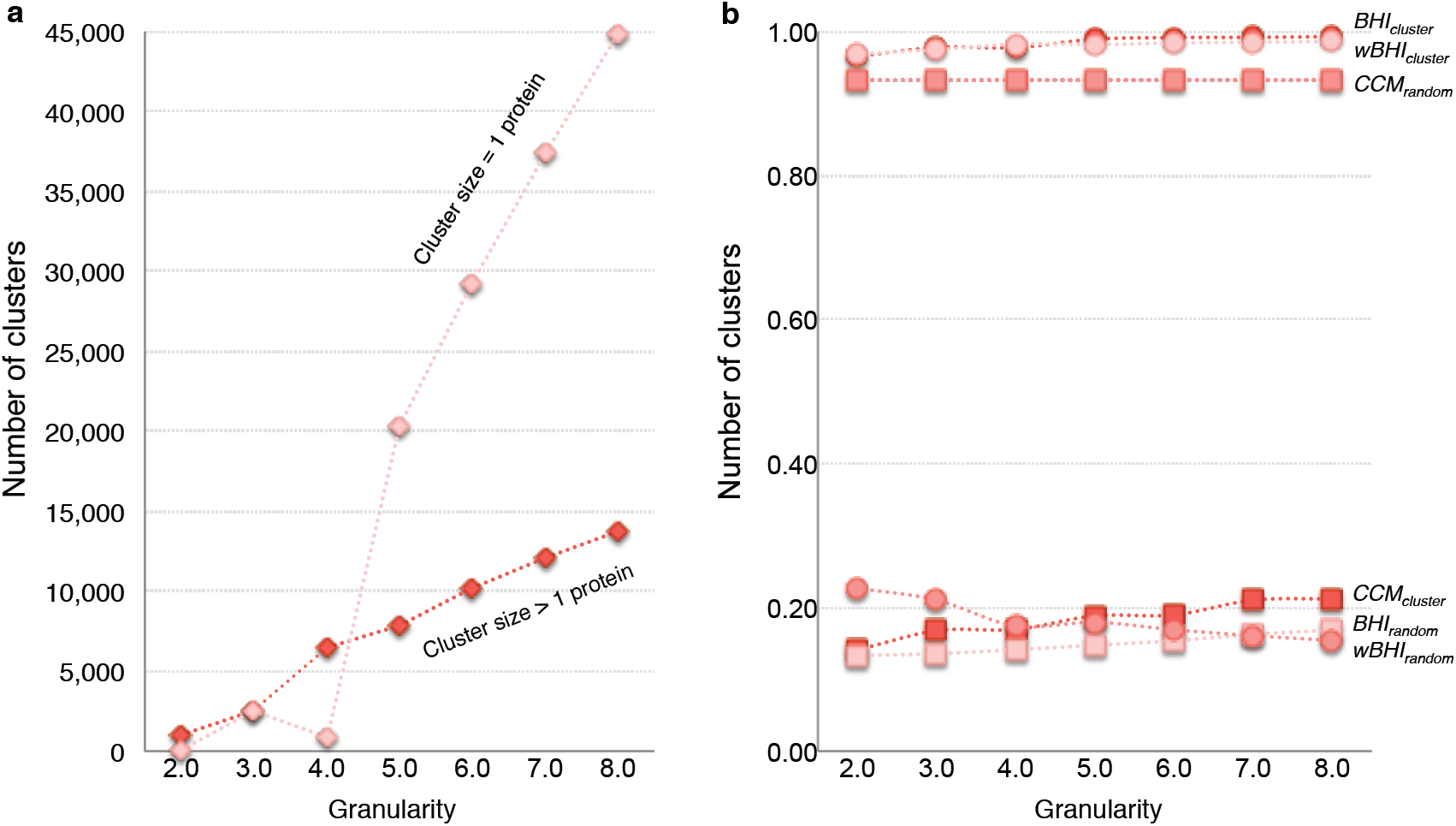
Influence of granularity on the clustering. (a) Number of clusters with more than one protein (dark diamonds) or clusters holding a single protein (pale diamonds). (b) *BHI* (dark), *wBHI* (pale) and *CCM* (medium) scores obtained with random clusters (squares) and normal clusters (circles), respectively.

### Assessment of the homogeneity of IS functional homologues

The homogeneity towards the functions of the proteins in the query set relied on the assumption that the first BLAST cut-off (10^-5^ *E*-value) was stringent enough to capture only functional homologues to the query proteins. Potential bias might nevertheless arise from query proteins possessing a supplementary functional domain unrelated to the IS role, or from the selection of proteins belonging to the same superfamily but differing in function. To address these issues, we calculated the functional vectors associated to each KEGG group or ACLAME family of the query set, as well as those for all obtained clusters. For a protein *P*_*i*_, we defined the associated functional vector with respect to its set of identified domains 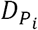. and to the set of all identified domains *D=*{*d*_*1*_,…,*d*_*x*_} as:

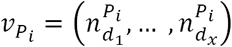

where 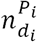 is the number of time *d*_*i*_ is found in 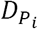. The functional vector associated to a given cluster of proteins *C*_*i*_ could then be defined as:

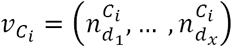

where 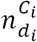 is defined as:

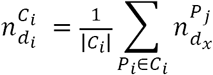

For each cluster *C*_*0*_, the cosine distance between its associated vector 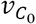 and the associated vector 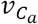 of the corresponding KEGG group or ACLAME family annotations *C*_*a*_ was then computed as:

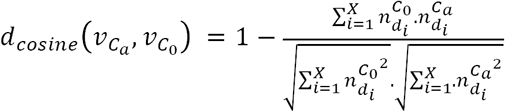

For each cluster *C*_*0*_, the cosine distance between its associated vector 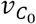 and the associated vector 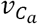 of the corresponding KEGG group or ACLAME family annotations *C*_*a*_ was then computed as:

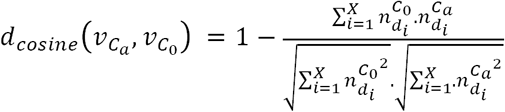

The 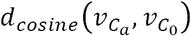 values were compared with those obtained using random clusters *C*_*r*_ of the same size than *C*_*0*_. For each *C*_*0*_ and its corresponding *C*_*a*_, 200 random clusters and their associated distances 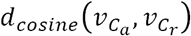, from which the corresponding empirical distribution *D*_*e*_ was constructed, were computed. *C*_*0*_ is then considered as noise if 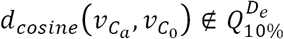 where 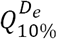 is the 0.1-quantile of *D*_*e*_.

### Unsupervised analyses of the replicon space

We represented the bacterial replicons (Supplementary Table 1) as vectors according to their content in IS genes. The number of IS protein clusters retained for the analysis determined the vector dimension and the number of proteins in a replicon assigned to each cluster gave the value of each vector component. We built matrices 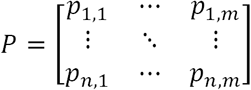, where *n* is the number of replicons, *m* the number of protein clusters, and *p*_*i,j*_ the number of proteins of the *j*^*th*^ cluster encoded by a gene present on the *i*^*th*^ replicon. We constructed several datasets to explore both the replicon type and the host taxonomy effects on the separation of the replicons in the analyses (Table 12).

**Table 12.** Reference classes used in the evaluation of the replicon IS protein-based unsupervised clustering solutions

| EVALUATED SEPARATION | ENSEMBLE | NORMALIZED ENSEMBLE<sup>a</sup> |
| --- | --- | --- |
| Chromosomes vs. Plasmids | $\{R^{\{chromosome\}}, R^{\{plasmid\}}\}$ | $\{\overline{K}l_{genus}^{R^{\{chromosome\}}}, \overline{K}l_{genus}^{R^{\{plasmid\}}}\}$ |
| Chromosomes *per* host phylum | $Kl_{phylum}^{chromosome}$ | $\overline{K}l_{genus}^K \vert K \in Kl_{phylum}^{chromosome}$ |
| Chromosomes *per* host class | $Kl_{class}^{chromosome}$ | $\overline{K}l_{genus}^K \vert K \in Kl_{class}^{chromosome}$ |
| Plasmids *per* host phylum | $Kl_{phylum}^{plasmid}$ | $\overline{K}l_{genus}^K \vert K \in Kl_{phylum}^{plasmid}$ |
| Plasmids *per* host class | $Kl_{class}^{plasmid}$ | $\overline{K}l_{genus}^K \vert K \in Kl_{class}^{plasmid}$ |

<sup>a</sup> Normalisation according to host genus

The taxonomic representation bias was taken into account by normalizing the data with regard to the host genus: a consensus vector was built for each bacterial genus present in the datasets. The value of each vector attribute was calculated as the mean of the attribute values in the vectors of the replicons that belong to the same bacterial genus. As a first approach, we transformed data into bipartite graphs whose vertices are the replicons and the proteins clusters. The graphs were spatialized using the force-directed layout algorithm ForceAtlas2 (Jacomy et al., 2014) implemented in Gephi (Bastian et al., 2009). Bipartite graphs are a powerful way of representing the data by naturally drawing the links between the replicons while enabling the detailed analysis of the IS cluster-based connections of each replicon by applying forces to each node with regard to its connecting edges. To investigate further the IS-based relationships of the replicons, we applied the community structure detection algorithm INFOMAP (Rosvall and Bergstrom, 2008) using the *igraph* python library (Csardi and Nepusz, 2006). We also performed a WARD hierarchical clustering (Johnson, 1967) after a dimension reduction of the data using a Principal Component Analysis (Hotelling, 1933). To select an optimal number of principal components, we relied on the measurements of the cluster stabilities using a *stability criterion* (Hennig, 2007) and retained the first 30 principal components (57% of the total variance). For consistency purpose, the number of clusters in the WARD analysis was chosen to match that obtained with the INFOMAP procedure. The number of clusters used was assessed by the stability index by Fang and Wang (2012) (Table 3). The quality of the projection and clustering results were confirmed using the V-measure indices (Rosenberg and Hirschberg, 2007) (*homogeneity, completeness, V-measure*) as external cluster evaluation measures (Table 3). The *homogeneity* indicates how uniform clusters are towards a class of reference. The *completeness* indicates whether reference classes are embedded within clusters. The *V-measure* is the harmonic mean between these two indices and indicates the quality of a clustering solution relative to the classes of reference. These three indices vary between 0 and 1, with values closest to 1 reflecting the good quality of the clustering solution. The type of replicons (*i.e.,* plasmid or chromosome) and the taxonomic affiliation (phylum or class) for chromosomes or plasmids were used as references classes (Table 12). Additionally, the *stability criterion* (Hennig, 2007) of individual clusters, weighted by their size, for a given clustering result was evaluated using the bootstrapping of the original dataset as re-sampling scheme. Individual Jaccard coefficient for each replicon were computed as the number of times that a given replicon of a cluster in a clustering solution is also present in the closest cluster in the resampled datasets.

### Functional characterization of the replicons and genomes

In order to characterize the functional bias of the replicons, 117 IS functionalities (46 from ACLAME and 71 from KEGG) were considered. When equivalent in plasmids and chromosomes, functions from ACLAME and KEGG databases were considered to be distinct. A *n*m* matrix 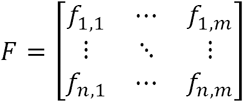 with *n* the number of replicons and *m* the number of IS functionalities, was used as input to the projection algorithms. *f*_*i,j*_ represents the number of times that genes coding for proteins annotated with the *j*^*th*^ function are present on the *i*^*th*^ replicon. Several datasets were analysed using PCA dimension reduction of the data followed by WARD hierarchical clustering (Table 3).

### Logistic regression analyses

Several reference classes of replicons and complete genomes were considered for comparison (Table 13). Ambiguous, *i.e.,* potentially adapted, plasmids belonging to INFOMAP clusters of plasmid replicons partially composed of SERs and/or chromosomes were removed from the plasmid class. When appropriate, the taxonomic representation bias was taken into account by normalizing the data with regard to the host genus as before. Logistic regressions (McCullagh and Nelder, 1989) were performed for the 117 IS functions using the R glm package coupled to the python binder rpy2. The computed *P*_*value*_ measured the probability of a functionality to be predictive of a given group of replicons/genomes and the *Odd-Ratio* estimated how the functionality occurrence influenced the belonging of a replicon/genome to a given group.

**Table 13.** Datasets used in the logistic regression analyses

| ENSEMBLE OF REPLICONS OR GENOMES | NOTATION | DATASET | DIMENSION<sup>a</sup> |
| --- | --- | --- | --- |
| Genus-normalized SERs | $E_{SER}$ | $\bar{V}_{f,genus}^{R\{SER\}}$ | (28, 117) |
| Genus-normalized plasmids | $E_{plasmid}$ | $\bar{V}_{f,genus}^{R\{plasmid\}}$ | (262, 117) |
| Genus-normalized chromosomes | $E_{chromosome}$ | $\bar{V}_{f,genus}^{R\{chromosome\}}$ | (560, 117) |

<sup>a</sup> (Number of replicons, number of functions)

### Supervised classification of replicons and genomes

In order to identify putative ill-defined SERs and chromosomes amongst plasmids, we performed supervised classification analyses using random forest procedures (Geurts et al., 2006). We used the IS functionalities as the set of features and the whole sets of chromosomes, plasmids and SER as sets of samples to build four classification studies (Table 7) and detect SER candidates (plasmids *vs.* SERs) and chromosome candidates (chromosomes *vs.* SERs or chromosomes *vs.* plasmids). Because of the unbalanced sizes of the training classes (SERs *vs.* chromosomes and plasmids), iterative sampling procedures were performed using 1000 random subsets of the largest class, with a size similar to that of the smallest class. The ensuing results were averaged to build the class probabilities and relative importance of the variables. We also used the whole set of plasmids when compared to SERs, to identify more robust SER candidates. The discarding of plasmids in the iterative procedure increases the classifier sensitivity while reducing the rate of false negatives by including more plasmid-annotated putative true SERs, whereas it decreases the classifier precision while increasing the rate of false positives. The ExtraTreeClassifier (a classifier similar to Random Forest) class from the Scikit-learn python library (Pedregosa et al., 2011) was used to perform the classifications, with *K=1000, max_feat=sqrt*(*number of variables*) and *min_split=1.* For each run, the *feature importances* and *estimate_proba* functions were used to compute, respectively, the relative contribution of the input variables and the class probabilities of replicons/genomes. The statistical probability of a replicon/genome belonging to a class was calculated as the average predicted class of the trees in the forest. The relative contribution of the input variables was estimated according to Breiman (2001). The choices of the number of trees in the forest *K*, the number of variables selected for each split *max_feat,* and the minimum number of samples required to split an internal node *min split* were cross-validated using a *Leave-One-Out* scheme. The performance of the *Extremely-randomized-trees* classification procedures was assessed using a stratified 10-fold cross-validation procedure following Han *et al.* (2012), and the out-of-bag estimate (OOB score) (Izzenman, 2008; Pedregosa et al., 2011) computed using the *oobscore* function of Scikit-learn python library.

## Data availability

The data supporting the findings of this study are available within the Article and its Supplementary Information or are available from the authors.

## SUPPLEMENTARY TABLES

Table 1. Replicon dataset

Table 2. INFOMAP IS protein-based clustering solution of the 4928 replicons

Table 3. PCA + WARD IS protein-based clustering solution of the 4928 replicons

Table 4. PCA + WARD IS function-based clustering solution of the 4928 replicons

